# SHIELD: Skull-shaped hemispheric implants enabling large-scale electrophysiology datasets in the mouse brain

**DOI:** 10.1101/2023.11.12.566771

**Authors:** Corbett Bennett, Ben Ouellette, Tamina Ramirez, Alex Cahoon, Hannah Cabasco, Yoni Browning, Anna Lakunina, Galen F. Lynch, Ethan McBride, Hannah Belski, Ryan Gillis, Conor Grasso, Robert Howard, Tye Johnson, Henry Loeffler, Heston Smith, David Sullivan, Allison Williford, Shiella Caldejon, Severine Durand, Samuel Gale, Alan Guthrie, Vivian Ha, Warren Han, Ben Hardcastle, Chris Mochizuki, Arjun Sridhar, Lucas Suarez, Jackie Swapp, Joshua Wilkes, Joshua H. Siegle, Colin Farrell, Peter A. Groblewski, Shawn R. Olsen

## Abstract

To understand the neural basis of behavior, it is essential to measure spiking dynamics across many interacting brain regions. While new technologies, such as Neuropixels probes, facilitate multi-regional recordings, significant surgical and procedural hurdles remain for these experiments to achieve their full potential. Here, we describe a novel 3D-printed cranial-replacement implant (SHIELD) enabling electrophysiological recordings from distributed areas of the mouse brain. This skull-shaped implant is designed with customizable insertion holes, allowing dozens of cortical and subcortical structures to be recorded in a single mouse using repeated multi-probe insertions over many days. We demonstrate the procedure’s high success rate, biocompatibility, lack of adverse effects on behavior, and compatibility with imaging and optogenetics. To showcase the scientific utility of the SHIELD implant, we use multi-probe recordings to reveal novel insights into how alpha rhythms organize spiking activity across visual and sensorimotor networks. Overall, this method enables powerful large-scale electrophysiological measurements for the study of distributed brain computation.

## Introduction

The neural mechanisms mediating behavioral and cognitive operations in the mammalian brain involve coordinated activity across highly distributed cortical and subcortical networks. Even simple sensorimotor transformations engage dozens of brain regions working in concert to convert sensory input into decisions and motor actions (Guo et al. 2014; Inagaki et al. 2022; Romo & Salinas 2003). The neural dynamics underlying these transformations have traditionally been studied one region at a time, but ongoing technical advances have opened new avenues for multi-regional measurements of neural activity (Machado et al. 2022). This is particularly true in the mouse, where new high-channel-count electrophysiology and large-scale calcium imaging have flourished. For instance, calcium imaging methods can survey activity across most of the dorsal cortex (W. E. Allen et al. 2019; Cardin et al. 2020; Musall et al. 2019; Ren & Komiyama 2021; Wekselblatt et al. 2016), including large field of view 2-photon microscopes that provide neuron-level resolution over millimeter scales (Kim et al. 2016; Sofroniew et al. 2016). However, calcium imaging is unable to capture single spikes and fast activity dynamics in neural networks, and subcortical brain regions are less accessible, typically requiring the removal of overlying brain tissue, or the implantation of a large diameter GRIN lens (Jung et al. 2004). In contrast, extracellular electrophysiology using multiple high-density probes, such as Neuropixels, provides the opportunity to record from distributed cortical and subcortical brain regions with high temporal fidelity and single spike resolution (Jun et al. 2017).

Multi-Neuropixels recordings have been used in mice to simultaneously record from 2-8 independently inserted probes (Chen et al. 2024; IBL et al. 2022; McBride et al. 2023; Peters et al. 2021; Siegle et al. 2021; Steinmetz et al. 2019; Stringer et al. 2019; Vesuna et al. 2020). These experiments are technically demanding, in part due to the challenge of accessing many brain regions in the same animal. Most previous studies have addressed this challenge by making multiple small craniotomies (*∼*1mm) or burr-holes through which individual probes are inserted to target regions of interest (Figure 1a). This approach has been used to record with 8 probes during spontaneous activity in awake mice (Stringer et al. 2019), 2-3 probes during a visual decision-making task (IBL et al. 2022; Steinmetz et al. 2019), 4 probes during dissociative drug administration (Vesuna et al. 2020), and up to 5 probes during a memory-guided movement task (Chen et al. 2024). Though creating multiple small craniotomies is a straightforward and flexible surgical procedure, there are several drawbacks to this method. First, the craniotomies are typically performed on the day of recording and require the mouse to be anesthetized. Mice are allowed to recover for a few hours before recording, but same-day anesthesia could alter natural neural dynamics and behavior (Bekhbat et al. 2016; Jacobsen et al. 2012; McKinney et al. 2022). Second, craniotomies can damage the underlying cortical tissue and, even in cases where no overt tissue damage occurs, can recruit a neuroinflammatory response that persists for several weeks (Holtmaat et al. 2009). Third, performing many craniotomies is time-consuming, and the risk of brain damage and adverse effects of prolonged anesthesia increases with the number of craniotomies, putting a practical limit on the number of regions that can be targeted in a single experiment. Lastly, although recordings are often performed from the same mouse for multiple days, each successive procedure increases the risk of infection and compromised brain health.

**Figure 1:**
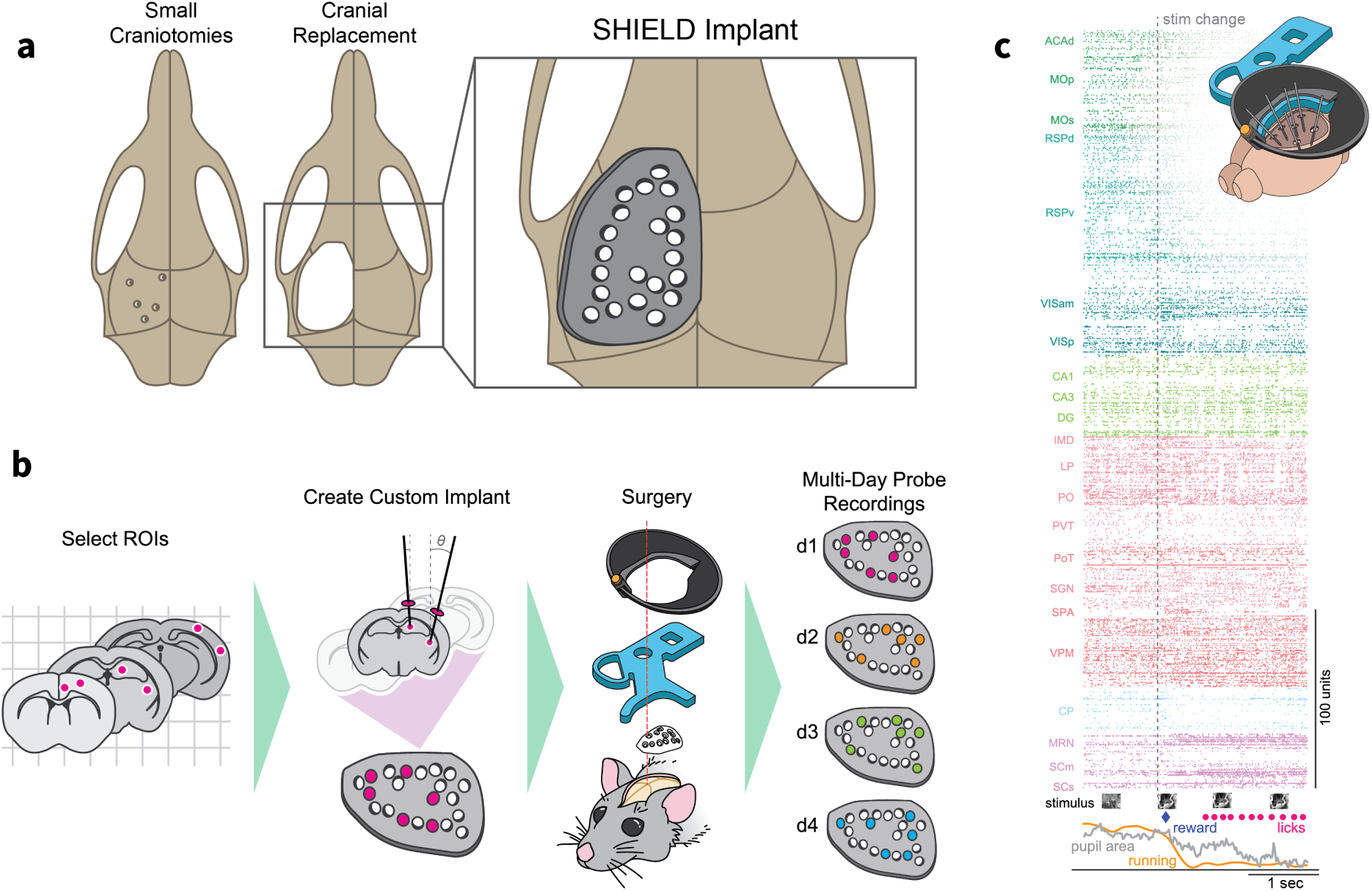
Overview of SHIELD implant and brain targeting strategy. **a,** Approaches for accessing the brain to insert multiple Neuropixels probes. **b,** Workflow for creating SHIELD implant to target distributed cortical and subcortical regions: stereotaxic targets are identified; rig geometry is used to calculate hole positions for each probe and an implant is fabricated; the implant is surgically implanted; multi-probe recordings are made over consecutive days. **c,** Raster plot showing spiking of 865 units across 22 areas in one example recording. During recording, the mouse performed a visual change detection task. Task-related behavioral data are aligned to a stimulus change (dotted line) after which the mouse licked to consume a water reward.

In our previous work, we developed an approach for accessing visual cortical regions of the mouse brain that employs an “insertion window” (Durand et al. 2023; Siegle et al. 2021). In this procedure, a large (5 mm) craniotomy is made over the primary visual cortex. To stabilize the brain, a plastic window with insertion holes for each probe is installed over the craniotomy before the recording. This technique enabled us to obtain high-quality, stable 6-probe recordings from precisely identified visual areas in hundreds of mice (Allen Institute 2023; Durand et al. 2023; Siegle et al. 2021). However, the procedure was developed for one or two acute recordings per animal, and probe insertions were limited to regions accessible through the cranial window.

To measure spiking activity across the mouse brain, we sought to develop a new procedure that would fulfill the following requirements: 1) provide flexible access to a broad set of brain areas, 2) enable simultaneous multi-probe recordings, 3) maintain brain health for many recording days, and 4) provide optical access to the brain for optogenetic perturbation and functional imaging to facilitate probe targeting.

To meet these requirements, we developed a novel cranial replacement implant and surgical workflow called SHIELD (<u>S</u>kull-shaped <u>H</u>emispheric Implants <u>E</u>nabling <u>L</u>arge-scale-electrophysiology <u>D</u>atasets) (Figure 1a). With this method, distributed brain regions are easily accessed for simultaneous silicon probe recordings via a 3D-printed SHIELD implant. The preparation allows flexible sets of multi-probe recordings to be made in a single mouse using repeated insertions over many days (Figure 1b). Moreover, the implant is transparent, allowing optical imaging, optogenetic perturbations, and optotagging to identify genetically defined subsets of neurons. Finally, we demonstrate the power of the SHIELD method during head-fixed behavioral experiments (Figure 1c) to reveal how alpha brain oscillations organize functional spiking interactions across distributed visual and sensorimotor networks.

## Results

### Design and 3D-printing of hemispheric skull implant

We sought to design a cranial implant that would fit mice within our standard surgical age range (postnatal day 56 ± 9; mean ± std). To conform the implant to the curvature of the mouse brain, we first performed cranial contour mapping in eight mice (Figure 2a). We then used CAD modeling to generate an ellipsoid shape that encompassed the set of skull contours (Figure 2a). This ellipsoid, defined by three curves with respect to bregma (Figure S1a), was easily parameterized for fast design iterations and was thus a convenient starting point for engineering the custom implants. To determine hole locations, stereotaxic targets (X, Y, and Z coordinates) were selected for each probe; the rig geometry was then taken into account to determine where each probe would need to enter the brain to reach its target (Figures 1b, 2b). Small insertion holes (0.75-1 mm) were then made in the implant CAD model at these locations (Figure S1c). In this study, we generated four distinct implants with 13-21 holes distributed broadly over the surface of the implant (Figure 2b and Figure S1e). In a smaller subset of mice, we also piloted a dual-hemisphere design (Figure 2c), which spans the midline and thus provides access to both the left and right hemispheres. To facilitate angled probe entry, the upper surface of each insertion hole was designed with symmetrical sloping cutaways (chamfers) (Figure 2b). To provide a surface that can be glued to the skull, a flange extends 600 µm beyond the cranial window. The inner part of the implant, circumscribed by this flange, extends 400 µm down into the craniotomy and sits on the brain surface (total thickness of implant: 650 µm; Figure S1a). Once complete, the implant CAD files are 3D-printed using clear resin at 25 µm layer height. After printing, the implants are washed in an isopropyl alcohol bath and then UV cured (Figure 2b).

**Figure 2:**
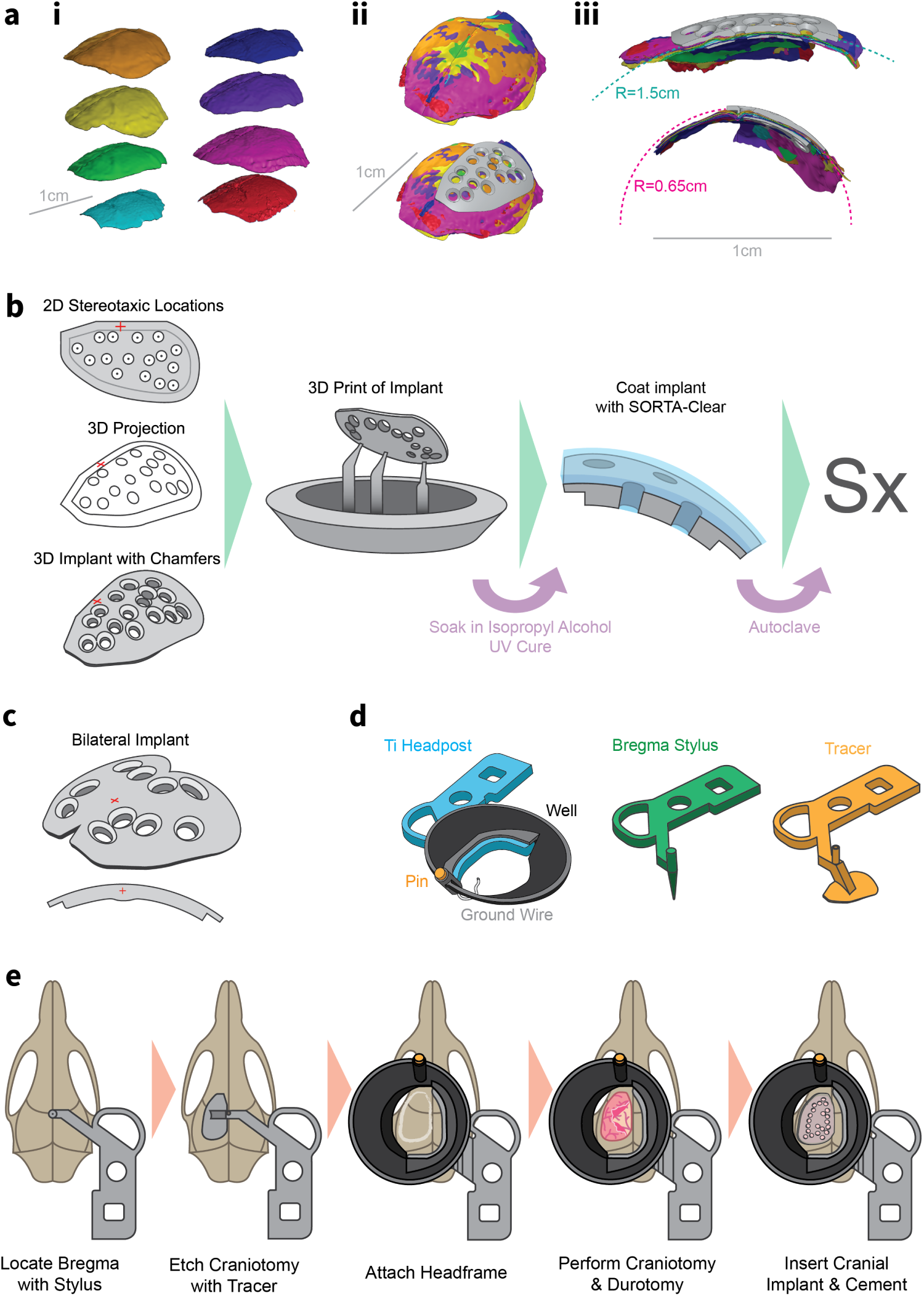
Design, preparation, and surgical installation of SHIELD implant. **a,** (i) Cranial contours obtained with laser scanning from eight mice, (ii) bregma-aligned composite contour with implant overlaid, and (iii) sagittal and coronal cross sections of implant overlays. **b,** Workflow for creating and preparing 3D-printed cranial implant. The set of desired insertion holes are defined in CAD software and chamfers are added. The implant is 3D-printed, cleaned, coated with SORTA-Clear, and then autoclaved. **c,** Example of a bilateral SHIELD implant that allows targeting regions in both left and right hemispheres. Cross-section at bregma shows the contoured ridge along the midline. **d,** Assembled headframe and custom tooling for locating bregma and tracing the outline of the craniotomy. **e**, Schematized view of the surgical workflow.

### Installation of SHIELD implant and quality control evaluation

Before surgically installing the cranial implant, SORTA-Clear 18 (Smooth-On) silicone rubber is applied to fill each insertion hole and form a thin layer on the implant’s dorsal surface (Figure 2b and Figure S1d). This layer of SORTA-Clear provides a solid substrate that plugs the holes prior to recordings, prevents deformation of the cortex by being flush with the implant, and maintains optical transparency necessary for regular brain health evaluations and through-implant imaging. The coated implants can then be sterilized by autoclaving.

To install the SHIELD implant, mice are anesthetized, placed in a stereotaxic frame, and the dorsal scalp removed. Bregma is located using tooling adapted from a previously described headframe and clamping system (Groblewski, Sullivan, et al. 2020) and serves as a reference point for placement of both the implant and the headframe. An outline of the implant location is etched into the skull using a custom tracing tool; then the headframe and well assembly is cemented in place (Figure 2d,e). A craniotomy is performed using the traced implant shape as a guide, and the prepared 3D-printed implant is placed in the opening of the craniotomy (Figure 2e). The edges of the implant are sealed to the skull using a light cure adhesive (Loctite 4305) and further reinforced with dental cement. Finally, a removable plastic cap is placed over the well to protect the implant’s silicone coating from cage debris.

To ensure the quality and reproducibility of the SHIELD method the following practices should be observed (Figure S2). Care should be taken when drilling near the veins and sinuses close to the implant location; in particular, the drill path comes near the rostral rhinal vein and dense vasculature around the sagittal sinus (Figure S2a). The implant has been designed to minimize drilling in this area, which can cause excessive bleeding. Over-drilling anywhere along the drill path can damage the underlying brain (Figure S2e). Generally, a very small but continuous crack in the final layer of skull is sufficient to achieve good separation. In addition, it is advised to keep a thin layer of bone along the lateral edge of the craniotomy (Figure S2b); this serves as a hinge and facilitates removal of the bone flap with fine forceps starting on the medial side (Figure S2c). To perform the durotomy, an incision is made along the anterior-posterior axis with a durotomy probe (Fine Science Tools), after which the dura is cut around the perimeter of the craniotomy using 45° Vannas scissors. The dura should be cleanly removed without tearing to avoid damaging underlying tissue or rupturing attached blood vessels along the midline (Figure S2f). The implant should be carefully aligned during placement using the anterior and medial edges of the craniotomy as reference points (Figure S2d). Improper seating of the implant into the craniotomy can lead to brain deformation (Figure S2g).

To validate the health of the brain following the SHIELD implant surgery, we performed a series of evaluations. First, we took images of the brain at multiple timepoints following the implant surgery. The transparent implant allowed visual inspection over eight weeks for abnormalities, including bleeding, tissue damage, bruising, and infection (Figure 3a). Of the 106 mice entering surgery for this study, 79 passed all quality checks with no issues, 14 passed with a flag that eventually resolved, and 13 mice failed (Figure 3b; Methods). The final passing rate of 87.7% is on par with the success rate for our previous 5 mm cranial window surgery used for multi-Neuropixels recordings (90%; 122/135 mice) (Durand et al. 2023; Siegle et al. 2021).

**Figure 3:**
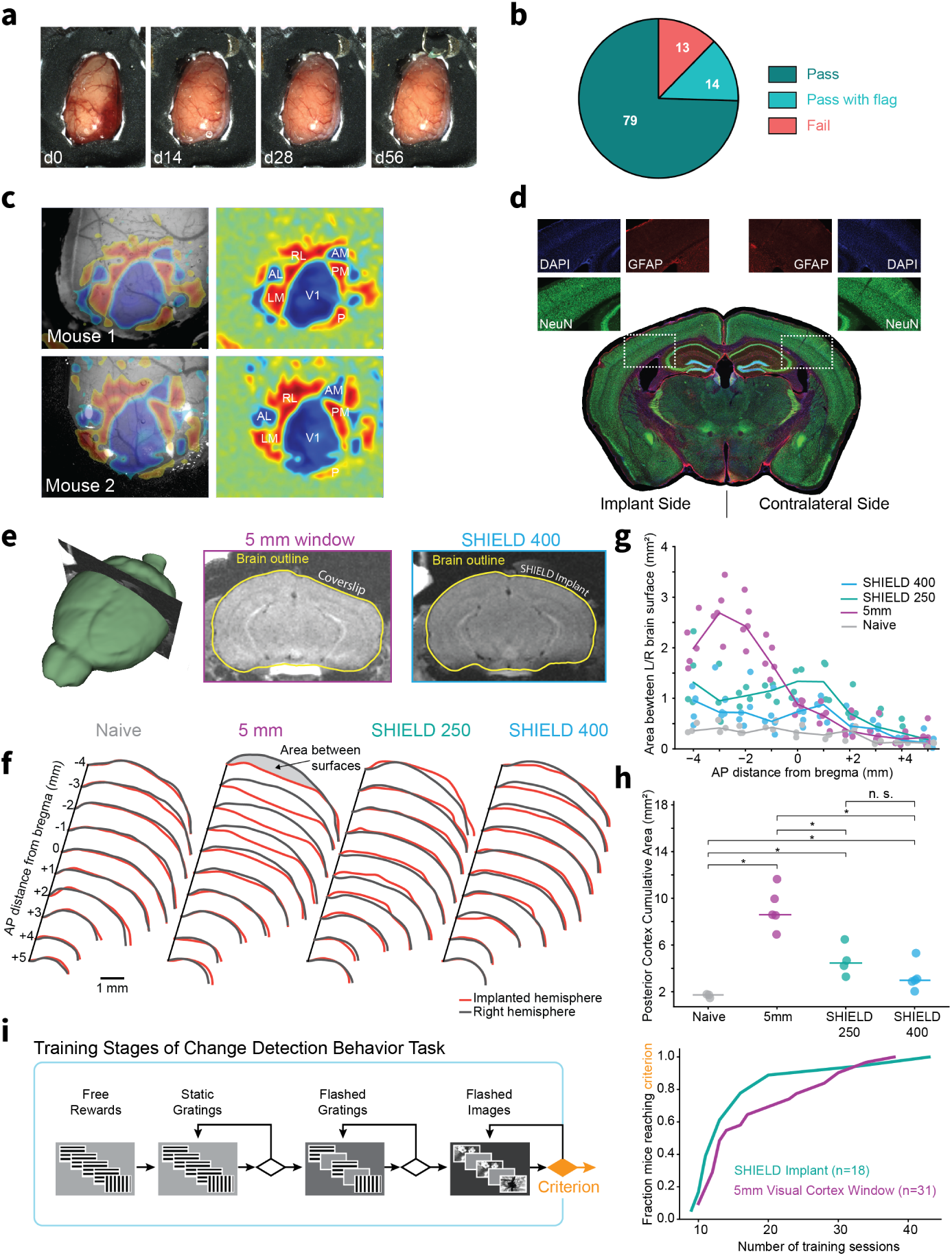
SHIELD procedure preserves health and function of underlying brain. **a,** Longitudinal images of implant in single mouse. **b,** Surgical outcomes from 106 implant surgeries in this study. **c,** Functional boundaries of the visual cortex mapped with through-implant intrinsic signal imaging. Left, visual sign map (Garrett et al. 2014) overlaid on implant. Right, sign map with visual area labels. **d,** Histology following implant surgery shows lack of neuronal loss (DAPI, NeuN) and absence of GFAP activation (also see Figure S3). **e,** Left: 3D MRI reconstruction of brain in mouse with SHIELD implant; Middle: Coronal slice from volume on left; Right: Coronal slice from MRI volume of mouse with 5 mm cranial window. **f,** Brain surface contours from MRI volume for naive (left), 5 mm window (middle), and SHIELD-implanted mice with the 250 µm and 400 µm insert designs (right). For comparison, the contour from the unimplanted hemisphere (gray) has been mirrored to overlay contour of implanted hemisphere (red). The area between contours was calculated to quantify brain deformation (gray shading). **g,** Area between surface contours for naive mice (gray, n=3), SHIELD-implanted mice (teal, SHIELD 250, n=4; blue, SHIELD 400, n=5) and mice with the 5 mm window (purple, n=5) along anterior/posterior (AP) axis. Lines indicate median across mice. **h,** Area between cortical contours summed across posterior AP sections ([-4 mm,-1 mm]; estimate of 5 mm window coverage) for each mouse. Line indicates median. * indicates p<0.05, Wilcoxon rank-sum test after Benjamini-Hochberg correction. **i,** Left: Training stages of visual change detection task. Mice progressed through four stages before reaching final behavior criterion (orange diamond). Right: Cumulative distribution of training sessions required to reach criterion for SHIELD (teal) and 5 mm window mice (purple).

Next, we characterized the functional integrity of the brain at the mesoscale level. We tested whether we could perform intrinsic signal imaging (ISI) through the implant to generate functional maps of the visual cortical areas using procedures previously developed for imaging through a glass window (Siegle et al. 2021; de Vries et al. 2020). We found that standard procedures successfully mapped the retinotopic boundaries of V1 and many higher visual cortical areas (Figure 3c). These results show that 1) the optical clarity of the cranial implant is sufficient for functional imaging, and 2) the functional properties of the visual cortical areas remain intact following the surgical procedure.

To determine if the implant caused a neuroinflammatory response or cell loss, we performed histology on the brains of implanted mice. We assayed for activation of glial fibrillary acidic protein (GFAP), a widely used marker of neuroinflammation (Eng & Ghirnikar 1994; Yang & Wang 2015), and examined cell density by staining with DAPI and NeuN. We compared GFAP, DAPI, and NeuN signal on the implanted left hemisphere to the unaltered right hemisphere and found little evidence of GFAP activation or cell loss, indicating that the implant did not compromise the health of underlying tissue (Figure 3d and Figure S3; n=6 mice, 56±12 days post-surgery; mean±stdev).

Because the implant is contoured to match the curvature of the skull, we hypothesized that the SHIELD procedure would minimize brain deformation in vivo compared with non-contoured implants. To visualize brain deformation in vivo, we performed magnetic resonance imaging (MRI) on mice with the SHIELD implant as well as mice implanted with a 5 mm cranial window centered on visual cortex (Figure 3e) and compared this data to publicly available MRI volumes from naive mice (Qiu et al. 2018). We extracted brain surface contours at ten coronal slices of the MRI volumes spanning the anterior-posterior extent of the dorsal cortex. Overlaying the surface contours for the implanted and contralateral control hemispheres revealed clear brain deformation for the non-contoured 5 mm window over visual cortex, evident as a separation between the experimental and control surface contours for posterior slices (Figure 3f,g). In contrast, the area between surface contours for SHIELD mice was more similar to naive mice (Figure 3f-h), indicating that the implant successfully replaced the skull without deforming the brain. This was true for two implant thicknesses we tested: one with a 250 µm thick cranial insert (500 µm total thickness; SHIELD 250) and one with a 400 µm insert (650 µm total thickness; SHIELD 400; see Methods).

Finally, we assessed whether behavior training times on a visual task were adversely impacted by the SHIELD implant. For this, we compared SHIELD implant mice to those with the 5 mm cranial window used in previous studies (Garrett et al. 2023; Groblewski, Ollerenshaw, et al. 2020). We found that SHIELD mice had similar learning rates to the 5 mm window mice (Figure 3i; median number of sessions to criterion: SHIELD 12.5±2; 5 mm window 14±3; p=0.17, Wilcoxon rank-sum test).

### High-quality multi-Neuropixels recordings

After successful installation of the SHIELD implant, mice in our study underwent behavioral training for 3-5 weeks before multi-probe recordings were performed. During training, the cranial implant remained covered with SORTA-Clear silicone plugs. Since Neuropixels probes cannot penetrate this durable silicone, it is removed prior to recording and replaced with a temporary Kwik Cast plug. On the recording day, this plug is removed, and the implant is covered with agar which fills the holes and provides mechanical stability (Figure 4a). The probes (each mounted on an independent micromanipulator) are then driven to their respective insertion holes and inserted through the agar into the brain (Figure 4a). Depending on the target structure, the probes are inserted to depths between 2-4 mm. Between daily experimental sessions (up to four consecutive recording days), the agar covering the implant is removed and replaced with a layer of Kwik Cast (Figure 4a). Alternatively, instead of sequential Kwik Cast and agar treatments, a *∼*1 mm thick layer of “Dura-Gel” (DOWSIL 3-4680) can be placed over the SHIELD implant (see Methods); this protects the brain but also permits Neuropixels probe insertions directly through the Dura-Gel layer. The Dura-Gel can remain in place for at least two weeks, allowing repeated probe insertions. Since Dura-Gel is not conductive, the ground wire must be placed beneath this layer, in contact with the brain surface.

**Figure 4:**
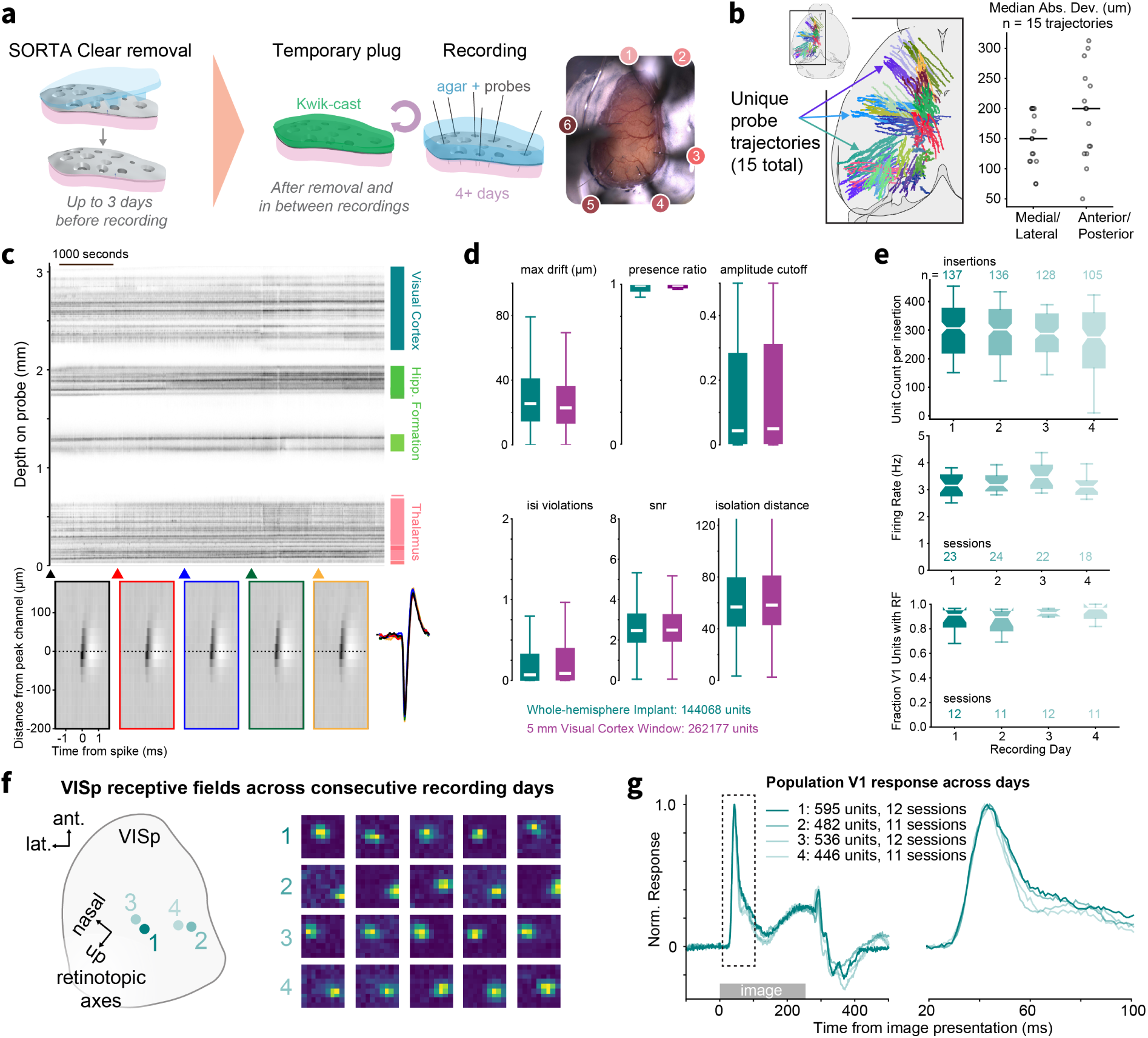
High-quality multi-Neuropixels recordings. **a,** Workflow for electrophysiological recordings. First, the SORTA-Clear covering that shielded the brain during behavior training is removed (up to 3 days prior to recording) and replaced with a temporary Kwik-cast plug. On the day of recording, this plug is removed and replaced with agar and a layer of silicone oil. After recording, the agar is removed and replaced with fresh Kwik-cast. **b,** (Left) Schematic of all probe insertion locations from this study registered to the CCF. Tracks are colored by insertion trajectory, defined by implant hole location and probe orientation. 15 unique insertion trajectories were examined in this study. (Right) The median absolute deviation in the medial-lateral and anterior-posterior axes across multiple insertions for each trajectory shown in (b). **c,** (Top) Drift map showing spiking activity across one probe during an example experiment. Darker dots indicate spikes with larger amplitude. (Bottom) Mean waveforms computed for five epochs across the recording duration for an example visual cortical unit to show waveform consistency. Arrowheads indicate the beginning of each epoch. The peak-channel waveform for all epochs are overlaid on the right. **d,** Unit quality metrics computed for this dataset (SHIELD implant) and the publicly available Allen Institute Neuropixels Visual Behavior dataset (5 mm window) for comparison. White band indicates median. Whiskers span the 10th and 90th percentiles. **e,** Functional unit properties across recording days. (Top) Unit count per probe insertion. (Middle) Overall session firing rate. (Bottom) Fraction of VISp units with significant receptive fields. Box plot notches indicate estimated 95% CI for median. **f,** Example receptive fields from one mouse across four consecutive recording days. Recording locations within VISp are diagrammed on the left. Axes indicate known retinotopic gradients for azimuth (nasal/temporal) and elevation (up/down). **g,** Mean population visual responses to one image across four consecutive recording days.

In a typical 6-probe experimental session, we recorded activity from over 1,800 units spanning 25 brain areas (this includes all non-noise units as described in Methods; 1821 ± 341.9 units; 28.1 ± 5.1 areas, mean ± std; Figure 4b). Across mice, insertions belonging to the same probe trajectory, defined by the probe orientation and the implant hole through which it was inserted, entered the brain at similar locations after post-hoc registration to the Allen Common Coordinate Framework (CCF) (n = 15 trajectories, median of the median absolute deviation (MAD) in CCF coordinates for each trajectory: 150 µm in the medial-lateral axis, 200 µm in the anterior-posterior axis; Figure 4b). Thus, our procedure allows users to consistently target areas of interest across animals with a dispersion of only 250 µm (a fraction of the 750 µm hole diameter).

A critical feature of extracellular electrophysiological recordings is the mechanical stability of the probe relative to the brain, since this can impact the fidelity of spike sorting and the ability to track neurons over the recording session—a factor that is particularly vital for recordings in behaving mice. In our experiments, mice were head-fixed but free to run on a disc while they performed a visual behavioral task, which required them to lick to report visual stimuli. We found that recordings were highly stable, as demonstrated by ‘drift maps’ that track the depth of each recorded unit on the probe over the duration of the experimental session (Figure 4c). We rarely observed discontinuous or abrupt shifts in the depth of units in the drift map (5.1% of insertions; 23/454). In addition, spike waveform shapes of individual units remained consistent over the course of the recording (Figure 4c). We quantified the drift of each unit in these experiments and found it was only slightly higher than that observed in our previous recordings using the 5 mm insertion window method (SHIELD: 25.1 ± 0.1 µm, 5 mm window: 22.82 ± 0.05; median ± sem; Figure 4d) (Durand et al. 2023; Siegle et al. 2021). We also note that drift can be reduced by retracting the probes 100 µm after insertion (Figure S4).

To further evaluate the recordings, we computed a battery of additional unit-level quality control (QC) metrics for experiments performed with the SHIELD implant as well as our previously used 5 mm visual cortex window recordings. This suite included metrics to evaluate unit stability and signal-to-noise (‘presence ratio’, ‘amplitude cutoff’, ‘snr’) as well as contamination from other units (‘isi violations’, ‘isolation distance’) (Siegle et al. 2021). Across all experiments, we found that the distribution of QC metrics was comparable between both surgical preparations (Figure 4d; median (bootstrapped 95% confidence interval): presence ratio: SHIELD 0.99 (0.99, 0.99), 5mm 0.99 (0.99, 0.99); amplitude cutoff: SHIELD 0.044 (0.043, 0.045), 5mm 0.049 (0.049, 0.05); ISI violations: SHIELD 0.07 (0.069, 0.071), 5mm 0.088 (0.088, 0.09); SNR: SHIELD 2.47 (2.47, 2.48), 5mm 2.5 (2.49, 2.5); isolation distance: SHIELD 56.1 (55.9, 56.3), 5mm 58.2 (58.1, 58.4); max drift: SHIELD 25.1 (24.9, 25.2), 5mm 22.8 (22.7, 22.9)). Indeed, we found that the percentage of units passing our standard quality metric filter (see Methods) was very similar for the SHIELD versus 5 mm window implants (% units passing quality metrics criteria: SHIELD: 41.1, 5 mm window: 40.7).

On each recording day, we measured the activity of nearly 300 units from each probe (Figure 4e). Unit yield was not significantly different across recording days (p=0.21 Kruskal-Wallis H-test). Additionally, the average firing rate of recorded units was similar across days (p=0.21 Kruskal-Wallis H-test), as was the fraction of VISp units that were significantly modulated by the receptive field-mapping stimulus (p=0.38 Kruskal-Wallis H-test). Together, these data provide additional confirmation that the brain remains healthy and stable over multiple days of consecutive recordings.

Next, we sought to validate the functional properties of neurons recorded using the SHIELD implant. To do this, we focused on visually-evoked activity in the primary visual cortex. In each of the four consecutive recording days, we measured many individual units with well-defined spatial receptive fields (Figure 4f ). On successive days we inserted the probe into a nearby location in the visual cortex using an adjacent hole in the implant; the region of visual space preferred on each day matched the expected location indicated by the ISI-based retinotopic map. In addition, the stimulus-evoked temporal dynamics were highly similar across the four recording days (Figure 4g).

The recordings described thus far were all performed with single hemisphere SHIELD implants. We also piloted recordings using the dual-hemisphere implant, demonstrating the feasibility of simultaneous, bilateral Neuropixels recordings with the SHIELD method (Figure S5).

### SHIELD implant is compatible with optogenetics in cortical and subcortical structures

Modern systems neuroscience experiments employ optogenetics to activate or silence components of neural circuits (Fenno et al. 2011), and to optotag genetic- or projection-defined cell types during in vivo recordings (Cohen et al. 2012; Lima et al. 2009). Thus, we tested whether we could optogenetically manipulate neural activity through the SHIELD implant.

First, we assessed cortical silencing by recording in VGAT-ChR2 mice (Zhao et al. 2011), which express channelrhodopsin in cortical inhibitory interneurons (Figure 5a). Light-activation of these interneurons produces potent network silencing with fast temporal dynamics (Guo et al. 2014; Li et al. 2019; Resulaj et al. 2018). Using a galvo-based laser delivery system, we shined blue light (488 nm) on the implant, systematically increasing the distance between the light and probe (0, 0.5, 1, 2 mm). During optogenetic stimulation we presented the animal with windowed grating stimuli (diameter: 50 degrees, duration: 250 ms) to verify that stimulus-evoked activity could be suppressed. The activity of regular spiking units (RS) in VISp was nearly silenced when the laser (2 mW power) was targeted to the recording location (Figure 5b). The effect of silencing was still strong 0.5 mm away but was similar to baseline 2 mm away (Figure 5c). The spatial extent of silencing varied with laser intensity (Figure 5c). This distance dependence matches what has been described for clear-skull preparations at similar laser powers (Li et al. 2019).

**Figure 5:**
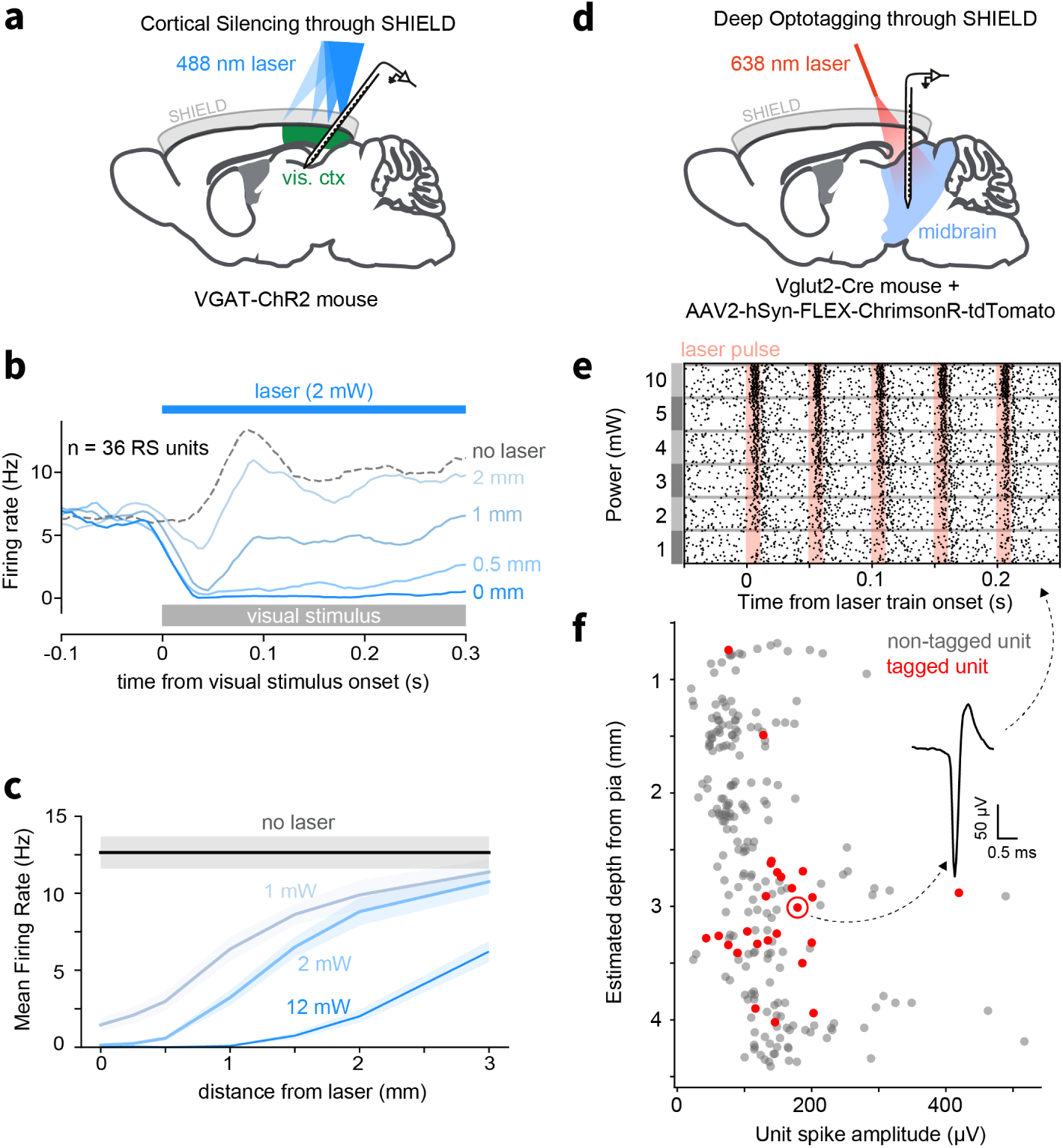
Optogenetics through SHIELD implant. **a,** Schematic of cortical silencing experiment with VGAT-ChR2 mouse. **b,** Efficacy of optogenetic silencing of cortex (VISp). Line intensity indicates the distance of the laser spot from the recording location. **c,** Spatial dependence of optogenetic silencing. **d,** Schematic of experiment for optotagging of VGLUT2+ neurons in the midbrain. **e,** Raster plot showing light-evoked spiking activity from an opto-tagged VGLUT2+ MRN neuron. Trials are grouped by laser power for display but were randomly interleaved during the experiment. **f,** Spatial distribution of opto-tagged and non-tagged units. The unit in e) is circled and its waveform shown in the inset.

Next, we tested whether illumination through the SHIELD implant would permit optotagging of neurons in deep subcortical structures. For these experiments, we recorded from the midbrain reticular nucleus (MRN) and sought to optotag glutamatergic neurons labeled by focal viral expression of the red-shifted opsin, ChrimsonR, in the midbrain of Vglut2-Cre mice (Figure 5d). Photo-stimulation with pulses (10 ms) of red light (638 nm) evoked reliable, short-latency spiking responses from units recorded in the midbrain (Figure 5e). Such optotagged units were found up to 4 mm below the surface of the brain (Figure 5f ).

Together, these experiments demonstrate that the SHIELD implant is compatible with targeted optogenetics combined with simultaneous Neuropixels recordings.

### Distributed recordings across cortical and subcortical circuits reveal distinct visual and sensorimotor modules during alpha-like oscillations

It has long been recognized that cortical activity, even in primary sensory areas, is profoundly modulated by the brain’s intrinsic dynamics (Buzsaki 2006). The dominant such brain rhythm in awake primate cortex is alpha activity (8-12 Hz), first documented by Hans Berger in the 1920’s (Berger 1929). Since this discovery, understanding how internally generated rhythms organize the flow of activity across brain regions remains a deep question in systems neuroscience.

Recent studies in mouse visual cortex have described low-frequency (3-5 Hz) oscillations in the membrane potential of superficial neurons (Arroyo et al. 2018; Einstein et al. 2017; Nestvogel & McCormick 2022) and in local field potential (LFP) recordings (Senzai et al. 2019). Like alpha activity, these oscillations occur during quiet wakefulness, and are abolished during periods of high arousal or locomotion (Arroyo et al. 2018; Nestvogel & McCormick 2022). They have therefore been proposed as an evolutionary precursor to the primate alpha rhythm (Senzai et al. 2019). Yet, since these studies were focused on primary visual cortex, it is unclear how these events impact brain-wide spiking activity.

We leveraged our SHIELD preparation to simultaneously monitor spiking activity across many cortical and subcortical brain regions. We then asked 1) when mouse alpha events occur in VISp, do they modulate spiking activity in other areas? If so, with what specificity? And 2) are alpha events in the mouse always associated with VISp activity, or can they engage distinct sets of cortical areas independently?

To answer these questions, we performed simultaneous recordings using up to six Neuropixels probes targeted to occipital, midline, and frontal cortical regions, as well as the hippocampus, thalamus, striatum, and midbrain. Consistent with previous studies, we observed prominent, rhythmic 3-5 Hz waves of activity in VISp (Figure 6a, left). These alpha events spanned all cortical layers and were characterized by brief spike bursts separated by longer silent periods lasting 200 ms, during which VISp neurons were almost completely inactive. These events were not restricted to VISp, but were commonly observed to coincide with synchronous bursts and silences across all visual cortical areas, as well as primary and secondary visual thalamus (LGd and LP), medial cortical structures (e.g. RSP), and the caudoputamen (CP).

**Figure 6:**
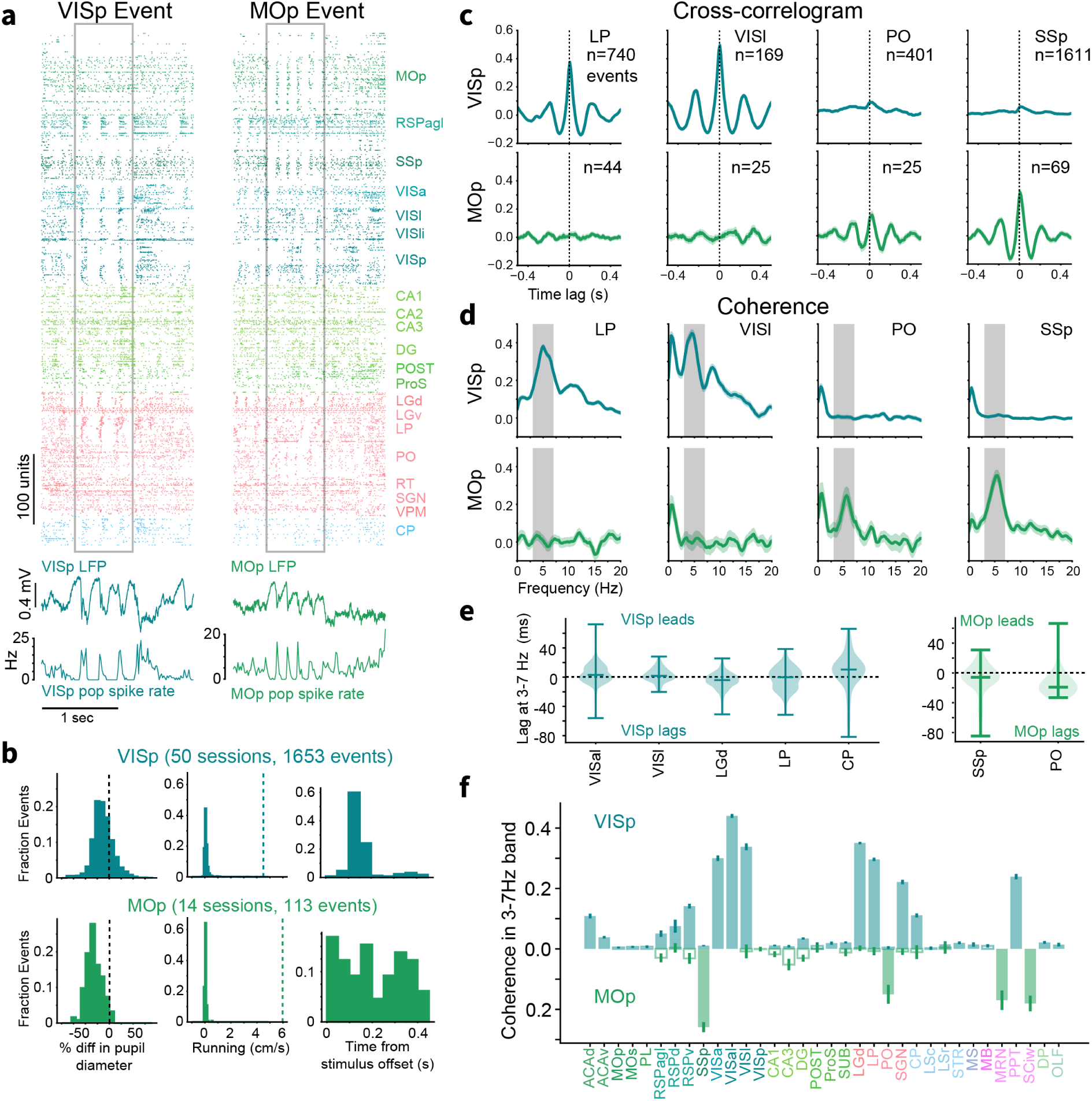
Distinct and distributed visual and motor networks during alpha-like cortical oscillations. **a,** Top: Population spike rasters triggered on example VISp (left) and MOp (right) oscillatory events. Both rasters are taken from the same recording session. Spikes are colored by their CCF area designation. Within a given area, units are ordered by depth. Bottom: LFP traces and population spike rates for VISp (left) and MOp (right) aligned to rasters above. **b,** (Left) Histogram of pupil diameter during alpha event relative to random experiment times for VISp events (top, blue) and MOp events (bottom, green). (Middle) Histogram of running speed during event relative to random times (dotted line). (Right) Histogram of event onset times relative to visual stimulus offset. **c,** (Top) Cross-correlograms of population spiking activity between VISp and LP, VISl, PO, and SSp during VISp events. (Bottom) As above, but between MOp and the same four areas during MOp events. **d,** (Top) Coherence in population spiking activity between VISp and LP, VISl, PO, and SSp during VISp events relative to random experiment times. Dark rectangle indicates 3-7 Hz band. (Bottom) As above, but for MOp during MOp events. **e,** Distribution of time lags in the 3-7 Hz band across during VISp-triggered (left) and MOp-triggered (right) events. **f,** Mean coherence in population spiking activity between VISp (top, blue) and 34 CCF brain regions during VISp events. Coherence between MOp (bottom, green) and the same 34 regions is plotted as downward going. Shaded boxes indicate significant values (p<0.05) after correction for multiple comparisons.

Notably, in the same recordings, we often observed similar rhythmic spiking in primary motor cortex (MOp) (Figure 6a, right). These MOp alpha-like events occurred in a slightly higher frequency band (5-7 Hz), and often coincided with synchronous activity in primary somatosensory cortex (SSp) and somatosensory thalamus (POm). Neither visual nor motor events appeared to engage the hippocampus.

To investigate these events more systematically, we algorithmically identified alpha activity based on rhythmic population spiking in either VISp or MOp (Methods). Like primate alpha rhythms, VISp and MOp events tended to occur during periods of low arousal when the animal was immobile and had a small pupil diameter (during alpha events, pupil diameter was 25% smaller relative to randomly chosen session times) (Figure 6b). The VISp events were often triggered by the offset of visual stimulation (Figure 6b), as described in previous studies (Arroyo et al. 2018; Einstein et al. 2017). Activity during post-stimulus alpha events was weakly but significantly correlated with activity during the preceding stimulus, indicating that alpha events tend to replay recent sensory stimulation (Figure S6; r=0.77, p=1.8e-6). The timing of MOp events was not related to visual stimuli (Figure 6b).

Initial inspection of example experiment rasters suggested that VISp and MOp events engaged distinct networks of cortical and subcortical regions (Figure 6a). To test this observation, we calculated cross-correlograms (CCGs) between population spiking activity in either VISp or MOp and other cortical and subcortical areas recorded during the alpha events (Figure 6c). During VISp events, VISp activity was tightly coupled to secondary visual thalamus (LP) and VISl (mouse V2), as evidenced by sharp peaks in the CCG at near-zero lag. VISp was not strongly coupled to somatosensory thalamus (PO) or primary somatosensory cortex (SSp). Indeed, coherence analysis revealed a prominent peak in the 3-7 Hz band between VISp population spiking activity and LP and VISl, but not POm or SSp (Figure 6d). Conversely, MOp showed the opposite pattern: strong coupling to PO and SSp in the 3-7 Hz band, but not to LP or VISl. Time lags between VISp and MOp and their associated regions in the 3-7 Hz band straddled zero lag, indicating these areas did not consistently lead or follow activity in the other network nodes (Figure 6e).

Quantifying coherence across all simultaneously recorded regions further revealed distinct visual and sensorimotor alpha rhythm networks (Fig 6f ). In addition to visual cortico-thalamic regions, visual alpha events enhanced coherence between VISp and medial cortical structures ACA and RSP, as well as the caudoputamen (CP), all of which are known to receive direct visual cortical projections (Oh et al. 2014). Conversely, MOp events enhanced coherence between MOp and SSp/POm, as well as motor-related midbrain structures MRN and deep layers of the superior colliculus (SCiw).

### Alpha-coherence reveals fine-scale functional coupling between visual cortex and striatum

We next asked whether alpha-coupling could reflect finer-scale functional connectivity between VISp and downstream targets. We chose to analyze the caudoputamen (CP) of the striatum, since it is known to receive direct projections from VISp and visual thalamus (Foster et al. 2021; Oh et al. 2014) and displays diverse sensory and motor-related responses (Peters et al. 2021; Reig & Silberberg 2014). We hypothesized that for a given insertion, CP units that displayed ‘pure’ visual responses to passively presented receptive field mapping stimuli (small gabor patches) would have stronger coherence with VISp during alpha events than CP units that did not have visual responses.

To test this, we first identified visually responsive CP units as those that were significantly modulated by the receptive field mapping stimulus (Figure 7a). This stimulus was shown outside the context of active behavior and was never associated with reward. We found that a population of CP units (CP-RF) did indeed display significant spatial receptive fields. These units were biased towards medial dorsal striatum mid-way along the anterior-posterior extent of CP (Figure 7b,c: note that we did not sample far lateral or posterior striatum).

**Figure 7:**
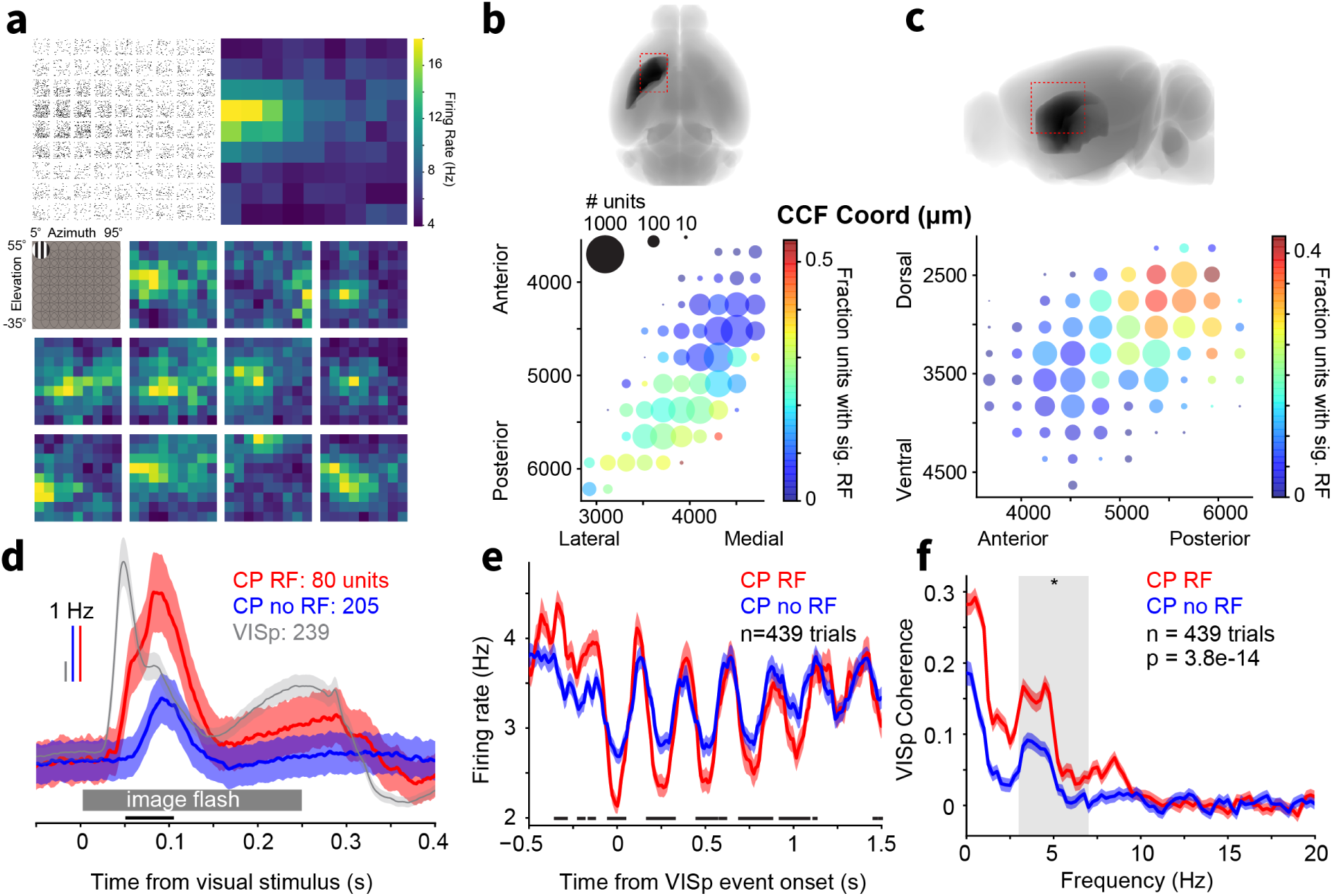
Visually-responsive striatal units are more tightly coupled to VISp alpha events. **a,** (Top) Receptive field measured for one example unit in CP displayed as a raster (left) showing spiking activity over 30 stimulus trials and as a heatmap of the mean response (right). (Bottom) Receptive fields for 11 more example CP units together with a diagram of the receptive field-mapping stimulus (upper left). **b,** (Top) Diagram of the mouse brain in horizontal orientation showing the position of CP (dark structure) and the CCF coordinates of the plot below (red dashed rectangle). (Bottom) Bubble plot depicting the recording locations of CP units projected onto the horizontal plane. The size of each circle indicates the number of units recorded at that CCF coordinate. The color indicates the fraction of units that had significant receptive fields. **c,** Same as B, but for the sagittal orientation. **d,** Population visual response of CP units with significant RFs (red), CP units without RFs (blue), and VISp units to a full-field natural image presentation for 250 ms. **e,** Population firing rates for CP-RF and CP-noRF units triggered on VISp alpha events. Black lines at the bottom of (d,e) indicate timepoints at which activity in the two CP populations was significantly different at p<0.05 after correction for multiple comparisons (Wilcoxon rank-sum test with Benjamini/Hochberg correction). **f,** Coherence of population spiking between the two CP populations and VISp during VISp alpha events.

Having split CP units into CP-RF and CP-noRF populations based on passive viewing of small gabor patches, we next compared their responses to full-field natural image presentations during active behavior. CP-RF units showed significantly larger visual responses to this stimulus compared to CP-noRF units (Figure 7d). In addition, the response time course of these two populations was different: CP-RF lagged behind VISp responses by 20 ms, whereas CP-noRF had slower responses that lagged behind CP-RF units by an additional 20 ms. This finding thus validates our parcellation of these units with an independent visual stimulus.

Finally, we tested whether CP-RF and CP-noRF units were differentially engaged by VISp alpha oscillations. We first computed population peristimulus time histograms (PSTHs) for the two CP groups triggered on the onset of VISp alpha events (Figure 7e). We found that CP-RF firing rates were significantly more modulated, particularly during the troughs of the oscillation when the visual thalamo-cortical network is suppressed. Next, we computed coherence between VISp and the two CP populations during VISp alpha events (Figure 7f ). As predicted, CP-RF units displayed higher coherence with VISp in the 3-7 Hz band, indicating that alpha events coordinate functionally related cortico-striatal subnetworks.

## Discussion

High-density silicon probes such as Neuropixels are transformative for systems neuroscience studies due to the nature and scale of the data they can collect. However, fully harnessing the power of these tools necessitates new surgical and experimental methodologies. In this study, we describe a highly flexible and robust workflow for engineering custom 3D-printed cranial implants that support user-defined multi-probe recording configurations across the mouse brain. This method delivers on the essential requirements for next-generation recordings in systems neuroscience: including brain-wide targeting, simultaneous multi-area recordings, and compatibility with behavior, optogenetics, and functional imaging.

Our method is highly robust and scalable: in this study we made stable, high-quality recordings from 25 mice, totaling 87 experiments and 513 probe insertions. Thus, this work goes beyond mere proof-of-concept and provides a rigorously vetted set of procedures for brain-wide recordings in behaving mice. The methods we describe should be easy to implement and adopt by many labs currently performing large-scale extracellular electrophysiological recordings with Neuropixels or other silicon probe devices. The SHIELD implant we describe is straightforward to fabricate in-house using a commercially available and relatively low-cost 3D printer. Moreover, we provide CAD files that can serve as a starting point for users to design their own custom implants using standard CAD software such as SolidWorks (https://portal.brain-map.org/explore/toolkit/hardware). Finally, because the same implant shape is consistent with many hole configurations, the surgical procedure can be standardized and practiced to improve consistency and raise success rates while allowing many different sets of brain regions to be targeted.

In a typical experiment for this study, we made simultaneous Neuropixels recordings from six probes that, in a single session, provided spike measurements from an average of 1,800 units across 25 brain regions. Moreover, an important aspect of the SHIELD preparation is that it supports multi-day probe insertions into different target regions. This expands the set of data that can be collected from each mouse and can reduce the number of mice needed for an experimental study. Multi-day recordings from the same mouse also provide powerful internal control data collected from a variety of brain regions (for instance, we were able to compare alpha-like rhythms from both visual and motor brain networks). Finally, multi-day recordings are important for experiments with behaving mice trained for weeks or months on complex behavioral tasks, where the value of each successfully trained mouse is considerable. In our experiments, we made recordings from behaving mice on four consecutive days. Thus, the SHIELD method provides high-yield, multi-area and multi-day recordings for large-scale spiking datasets in the context of active behavior.

### Comparison with other methods

Most previous studies making multi-Neuropixels recordings in the mouse brain have used the multiple small craniotomy approach in which holes are drilled into the mouse skull on the day of the recording (Figure 1a) (Chen et al. 2024; IBL et al. 2022; McBride et al. 2023; Steinmetz et al. 2019; Stringer et al. 2019; Vesuna et al. 2020). As described in the Introduction, this approach, while flexible, has many drawbacks. Thus, the method we describe here should prove extremely valuable for future studies that seek to make multi-probe recordings from distributed regions of the mouse brain.

Our cranial-replacement approach was inspired by prior methods for cortex-wide optical imaging in which a large fraction of the mouse skull is replaced with a transparent window (W. E. Allen et al. 2017; Ghanbari et al. 2019; Kim et al. 2016; Wekselblatt et al. 2016). For instance, the “See-Shells” method is a multi-component cranial implant assembly that includes a PET film that directly covers the dorsal cortex after skull removal surgery (Ghanbari et al. 2019). The PET film was optimized for optical clarity to support functional imaging, but Ghanbari et al. also made proof-of-concept electrophysiological recordings through a single perforation in the PET film, made after the implant was installed on the brain. In our study, we generated SHIELD implants with up to 21 pre-designed holes; this allows many distinct multi-probe insertion configurations, each targeting different sets of brain areas for simultaneous recording. Moreover, our procedure simplifies the experimental preparation, avoiding anesthesia and mitigating the risk of brain damage before recordings.

More generally, previous studies have designed 3D-printable skull-attached assemblies (including probe holders and head-caps) that support targeting and implantation of multiple electrode arrays or tetrodes to record from distributed brain regions in the rat (L. M. Allen et al. 2020; Headley et al. 2015; Ma et al. 2019; Sheng et al. 2021; Vöröslakos et al. 2021) and mouse (Chung et al. 2017). Conceptually these implants differ from the SHIELD implant. First, the SHIELD implant physically replaces the skull of the mouse, with customizable insertions holes providing brain access; conversely, the skull-attached assemblies are affixed to the intact cranium and still require individual craniotomies to be performed over each point of insertion. Second, the SHIELD implant is designed to support repeated acute multi-probe recordings, enabling researchers to target different sets of brain areas in the same animal over multiple experimental sessions. In contrast, the 3D printed skull-attached assemblies are tailored for chronically implanted electrode arrays that remain fixed in place or have linear travel via a microdrive.

### Future developments

The majority of experiments described in this study used single-hemisphere SHIELD implants providing access to the left hemisphere of the mouse brain. Many experimental applications, including those involving bilateral sensory stimulation and spatially directed motor control, necessitate recordings from brain regions across both hemispheres. To support this, we piloted a bilateral SHIELD implant. The larger implant requires a more complicated surgery in addition to headframe modifications. However, our experiments demonstrate the feasibility of this method for such bilateral recordings (Figure S5). Future work will continue to optimize the dual-hemisphere implant to expand the range of targets and improve the surgical procedure.

Because the implant is transparent, it can be combined with optical imaging to map the functional boundaries of cortical areas, as we demonstrate (Figure 3c). Importantly, the implant also provides the possibility for simultaneous multi-modal imaging, optogenetics, and distributed electrophysiology. Future studies could use SHIELD implants for paired 1-photon widefield calcium imaging plus Neuropixels recordings; imaging would measure broad activity dynamics across the cortex, while multi-Neuropixels would measure spiking activity from distributed cortical and subcortical regions (Peters et al. 2021).

The recordings we describe in this study involve acute insertions of multiple Neuropixels probes at the start of the recording session and removal at the end. Such acute recordings have proven extremely valuable for studying neural activity during sensory processing, decision-making, and behavior. It is also critical to measure how activity dynamics in the brain evolve over the course of days, weeks, and months. Previous studies have demonstrated the feasibility of chronic multi-channel electrode recordings in mice (van Daal et al. 2021; Juavinett et al. 2019; Steinmetz et al. 2021). The implant we describe here could be combined with skull-attached assemblies that support such chronic recordings (L. M. Allen et al. 2020; Chung et al. 2017; Headley et al. 2015; Ma et al. 2019; Vöröslakos et al. 2021).

### Example scientific use case: alpha oscillatory events engage distinct cortical-subcortical networks

Alpha rhythms are the strongest electrophysiological signature of the awake brain and have been the subject of intense study since their first description by Hans Berger almost a century ago (Berger 1929). Originally thought to reflect cortical ‘idling’ (Pfurtscheller et al. 1996) due to their prominence during quiet wakefulness, they have since been hypothesized to play an important role in regulating information transmission between cortical areas to subserve a number of cognitive processes, including working memory and attention (Saalmann et al. 2012; Von Stein & Sarnthein 2000). Human studies have typically measured alpha activity with non-invasive electrophysiological techniques such as EEG and MEG or, in some cases, intracranial electrocorticogram arrays (Halgren et al. 2019). In non-human primates, laminar probe arrays have measured alpha rhythms across the depth of cortex in multiple sensory regions (Haegens et al. 2015). However, interpretation of these results is complicated by poor spatial resolution and volume conduction from nearby structures. Moreover, these techniques are indirect measures of neural activity and thus cannot reveal the spike output of a given region, much less the activity of individual neurons, making it impossible to determine whether alpha events engage circuits with cell-type specificity.

Together with recent advances in silicon probe technology, our preparation is ideally suited to capture coordinated activity across distributed cortical and subcortical regions with neuron resolution. We therefore set out to characterize how alpha events shape spiking activity across diverse brain areas. Building on recent studies that have described alpha-like oscillations in mouse primary visual cortex (Arroyo et al. 2018; Einstein et al. 2017; Nestvogel & McCormick 2022; Senzai et al. 2019), we extended these findings in several important ways.

First, we show that alpha events identified in VISp reach beyond the VISp-LGd cortico-thalamic loop, modulating spiking activity across higher visual areas but also medial cortical regions RSP and ACA, as well as the striatum. Taken together with our observation that alpha events in VISp replay recent sensory activation (Figure S6), our data support a role for alpha rhythms in facilitating plasticity across the wider visual network in the mouse brain (Han et al. 2008).

Second, we find that alpha events in the mouse are not restricted to the visual cortex. We recorded similar rhythmic patterns of spiking activity in MOp. Strikingly, MOp and VISp events engage complementary networks of cortical and subcortical brain regions, with MOp showing tight coupling to SSp and secondary sensory thalamic nucleus POm and midbrain motor structures. Like VISp alpha activity, MOp events occurred during periods of low arousal and immobility and thus resembled a motor-analog of alpha in primates, typically referred to as the mu rhythm (Arroyo et al. 1993). In rats, mu-like events (termed high-voltage spindles) have been studied in the sensorimotor system (Buzsaki et al. 1988) and found to propagate to the brainstem (Nicolelis et al. 1995). These findings suggest that alpha activity should be construed as a family of rhythms (Buzsaki 2006), potentially generated by similar mechanisms but coordinating activity across distinct subnetworks of functionally related cortical and subcortical regions.

Finally, we leveraged the single-neuron resolution of Neuropixels recordings to investigate fine-scale functional interactions between VISp and the striatum during alpha events. We found a population of CP units with robust visual receptive fields. Though visual responses in mouse striatum have not been extensively characterized, the CP-RF cells described here resemble neurons recorded in the primate caudate (Hikosaka et al. 1989), which respond to visual stimuli unconditional on motor activity and have large, long-latency spatiotemporal receptive fields. CP-RF cells were intermixed with CP-noRF cells but were biased towards intermediate dorsomedial regions of the striatum known to receive visual projections (Bennett et al. 2019; Hintiryan et al. 2016). During VISp alpha events, CP-RF cells had greater firing rate modulation and higher coherence with VISp population activity than CP-noRF cells, suggesting fine-scale functional specificity in VISp-striatal coupling. In rats, motor-related dorsolateral striatum participates in high-voltage spindles (Berke et al. 2004), and coherence between striatal cells and MOp was found to increase over the course of learning a brain-machine interface (BMI) task (Koralek et al. 2013). In that task, cortico-striatal coherence was specific to the population of MOp neurons used to control the BMI. Together with our findings, these results suggest that highly specific coupling during alpha activity is a hallmark of cortico-striatal interactions. Such specificity may play an important role in plasticity and learning.

## Conclusion

To understand the intricate neuronal processes underlying complex behavior and cognition, it is imperative to simultaneously measure fast timescale spiking activity across numerous interacting brain regions. While advancements like Neuropixels have enabled increasingly large numbers of neurons to be recorded, substantial methodological and surgical challenges remain to fully capitalize on their potential, particularly for concurrent multi-probe recordings. Our innovative SHIELD method provides an elegant solution for accessing and recording from the mouse brain with many simultaneously inserted Neuropixels probes, allowing dozens of regions and thousands of neurons to be measured in a single mouse. This timely addition to the field enables cutting-edge experiments that promise to uncover new insights into the mechanisms of distributed computation and multi-regional signal flow dynamics across the brain.

## Materials and methods

### Mice

Mice were maintained in the Allen Institute animal facility and used in accordance with protocols approved by the Allen Institute’s Institutional Animal Care and Use Committee. Electrophysiology experiments used C57BL/6J wild-type mice purchased from Jackson Laboratories at age P25-50 (n = 7) and four transgenic lines (n = 1 VGAT-ChR2, n = 14 Sst-IRES-Cre; Ai32, and n = 4 Vip-IRES-Cre; Ai32, n=1 Vglut2-IRES-Cre;C57BL/6J, 18 males and 9 females). For skull contour mapping, eight mice were used (n= 3 Vip-IRES-Cre;Ai148, n=4 Slc17a7-IRES2-Cre;Camk2a-tTA;Ai93, n=1 Sst-IRES-Cre;Ai148, 5 males and 3 females). For MRI, 14 mice were used (5 mm cranial window: n=4 C57BL/6J, n=1 Vip-IRES-Cre;Ai14, 5 males; SHIELD 400 implant: n=5 C57BL/6J, 3 males and 2 females; SHIELD 250 implant: n=2 C57BL/6J, n=1 Ai230;C57BL/6J, n=1 Ai229;C57BL/6J, 4 males). Following surgery, all mice were single-housed and maintained on a reverse 12-hour light cycle in a shared facility with room temperatures between 68° and 72°F and humidity between 30 and 70%. All experiments were performed during the dark cycle. During behavior training, mice were water restricted to 85% of their initial body weight, with ad libitum access to food.

### Headframe

To enable co-registration across surgical, intrinsic signal imaging, and electrophysiology rigs, each animal was implanted with a 3D-printed, grade 5 titanium headframe (Groblewski, Sullivan, et al. 2020). In addition to a standard clamping interface, the headframe includes a modified L-shaped portion that contacts the right hemisphere parietal skull plate. The headframe is affixed with a black acrylic photopolymer well with an exposed gold pin for grounding recordings. For MRI experiments, a headframe with a slightly modified design was milled out of zirconia (Top Seiko).

### Implant design

The implant dimensions are defined in CAD. For the single-hemisphere implants used in our electrophysiology experiments, the total thickness of the implant was nominally 500 µm, with the ‘stump’ portion (i.e., the part that is inserted into the craniotomy) 250 µm thick, and the ‘flange’ 250µm thick. After MRI evaluation, we revised the design to include a slightly thicker stump (400 µm for a total thickness of 650 µm), which we found to better mimic the interior surface of the skull. The flange is larger than the craniotomy by 600µm and sits on top of the mouse skull, which provides an area to apply glue and to set the implant orientation in 3D space.

To design the implant in CAD software, we started with a triaxial ellipsoid-like surface matching the curvature of an average mouse skull. This design was defined by three curves positioned with respect to Bregma (Figure S1). The ellipsoid-like surface is then generated using the Solidworks ‘Surface Fill’ function with the patch boundary being the ellipse, and the constraint curves being the circular section curves. The next step is to cut the ellipsoid-like surface, once for the shape of the craniotomy, and again for the shape of the flange, resulting in two surfaces. The final shapes for these surfaces were created using an iterative feedback process, during which surgeons tested prototypes and provided feedback referenced to stereotaxic measurements. Once the craniotomy and flange surfaces have been cut, holes are sketched on the Bregma-Lambda plane and dimensioned with respect to Bregma. A typical hole size is 750 µm. No holes should be in the flange portion of the implant, and holes should not overlap. Holes are cut into both the stump and flange surfaces using the ‘surface-trim’ feature in Solidworks. ‘Thicken’ operations are performed on the two surfaces. The larger flange surface is thickened up 250 µm. The smaller stump surface is thickened down 400 µm. The final step of the design process is to add 45-degree chamfers to the tops of the insertion holes. The chamfers allow flexibility in probe angles and also make it easier to fill the holes with SORTA-Clear.

The dual hemisphere design and creation process is similar to the single hemisphere design. The perimeter was changed to include the left hemisphere and to avoid sagittal confluences in the anterior and posterior regions. We found that simply extending the single hemisphere implant led to gaps over the midline portion of the skull and thus needed to adjust the geometry of the underlying triaxial ellipsoid-like surface. Lastly, we added a protrusion on the underside of the implant along the midline to match the brain’s groove between the two hemispheres. The method of cutting a craniotomy perimeter out of a triaxial ellipsoid-like surface, thickening, and then cutting insertion holes to form the dual hemisphere implant was otherwise identical to the single hemisphere implant.

Implant CAD files are exported as .stl files and imported into Formlabs PreForm for printing on a Formlabs Form 3+ printer in clear resin at 25 µm layer height. After printing, the implants are washed in a dedicated isopropyl alcohol bath (Formlabs Wash Station) maintained at 99% concentration for 10 minutes. The implants are then UV cured in the Formlabs Cure station at 60°C for 15 minutes. Prior to implanting, the implant is coated with SORTA-Clear 18 silicone rubber (Smooth-On; Figure 2a). SORTA-Clear improves optical clarity for brain health evaluations and optical imaging and protects the brain until electrophysiology recordings. SORTA-Clear is mixed and degassed following manufacturer recommendations. Implants are pressed into a putty mold (ComposiMold, ImPRESSive putty) to seal the underside from leaks while the SORTA-Clear cures (Figure S1). The SORTA-Clear is drawn into a 1mL syringe, then a blunt 25G needle is attached for dispensing into the holes. Each hole must be filled individually and slowly to prevent trapping air. After each hole is filled, a thin layer is applied over the dorsal surface of the implant overlying the craniotomy, and the molds are left to cure for 24 hours. Coated implants are then cleaned by rinsing in 70% ethanol, placed into individual autoclave bags, and autoclaved for 5 minutes (131°C, 14.7 psi). Note that autoclaving at higher temperatures can lead to defects in the implant.

### Surgical procedure for hemispheric craniotomy

A pre-operative injection of dexamethasone (3.2 mg/kg, S.C.) is administered 1 h before surgery to reduce swelling and postoperative pain. Mice are initially anesthetized with 5% isoflurane (1-3 min) and placed in a stereotaxic frame (Model# 1900, Kopf ). Isoflurane levels are maintained at 1.0-2.0% and body temperature is maintained at 37.5°C for the duration of the surgery. Carprofen is administered for pain management (5-10 mg/kg, S.C.), and atropine is administered to suppress bronchial secretions and regulate heart rhythms (0.02-0.05 mg/kg, S.C.). An incision is made on the dorsal surface of the skull, and skin is removed in a teardrop shape, exposing the rostral rhinal vein between the eyes, the dorsal surface of the parietal and occipital skull plates, and stopping where the neck muscle begins to attach to the back of the skull. Next, the periosteum is removed from the skull surface to improve adhesion of the cement to the skull and prevent future scabbing and infection. Starting posterior of the left eye, angled forceps are used to separate cheek muscle from the skull, as well as connective tissue and muscle above the left ear. The cheek muscle is then pulled away from the skull and is stretched out so that it makes a seal with the left lateral portion of the well. The exposed skin (and any exposed soft tissue such as the cheek muscle) is then sealed with Vetbond, and the exposed skull is leveled with respect to pitch, roll, and yaw. Once the skull is level, bregma is identified using the custom bregma stylus (Figure 2c). Without moving the stereotaxic arm in X or Y, the stylus is replaced with a custom “tracer” (Figure 2c) that provides a guide for marking the craniotomy with respect to bregma. A #11 scalpel blade (or forceps) is used to etch a faint line in the skull around the tracer, which is then replaced with a shallow trench by lightly drilling, without breaking through the skull (NeoBurr EF4). After etching is complete, the tracer is replaced with the headframe, which is then lowered in Z to make contact with the skull. Next, dental cement (C&B Metabond, Parkell) is used to attach the headframe to the skull. Once the cement has hardened, the headframe is clamped into a custom frame and the craniotomy and durotomy are performed. The autoclaved custom implant is then attached to the skull by applying light-curable glue (Loctite 4305) between the flange and skull which is then hardened with blue light. Any areas of exposed skull are then covered with cement. Any white cement on the inside of the well is coated with a layer of black cement to reduce glare during ISI imaging. The mouse is given 1.0-1.5mL Lactated Ringers Solution (LRS) to help recover from the surgery and replace lost fluids. After removing the mouse from anesthesia, but prior to it waking up, a spatially registered image of the cranial window is obtained using a custom imaging system. Finally, a removable plastic cap is placed over the well to protect the coated implant from cage debris, and the mouse is returned to its home cage for recovery. Over the following 7-14 days, mice are monitored regularly for overall health, cranial window clarity, and brain health. Mice were failed out of the experimental workflow for detached headpost (4/106), reoccurring seizures (1/106), issues with the implant (2/106), brain health issues (1/106), or unexpected death (3/106). Other issues were flagged in 14/106 mice but were successfully resolved; these included scabbing, dehydration, transient seizure, and anal prolapse.

In a subset of mice, we modified the above procedure to perform a larger dual hemispheric craniotomy with the substitution of a suitable headframe, tracer, and implant. Here we note the key procedural differences. First, the thick bone overlying the sagittal sinus was drilled very carefully to achieve complete separation from the adjacent skull. Failure to do so often led to tearing of the sinus and excessive bleeding. Next, the left and right anterior-most sections of the drill path were partially drilled (though again taking care to completely separate the V shaped notch that crosses the sinus in between). This is to provide stability when removing the large piece of skull formed by the drill path. To remove the island of bone formed after drilling, the skull flap was gripped from the posterior edge to minimize tearing of attachment points along the sagittal sinus. Finally, the durotomy was performed in two stages—one for each hemisphere, as the durotomy cannot extend beyond the sagittal sinus.

### Intrinsic signal imaging

Intrinsic signal imaging (ISI) was used to delineate functionally defined visual area boundaries (Garrett et al. 2014; Juavinett et al. 2017) to enable targeting of Neuropixels probes to retinotopically defined locations in primary and secondary visual areas as described previously (Siegle et al. 2021). Briefly, mice were lightly anesthetized (1-1.4% isoflurane) and eye drops (Lacri99 Lube Lubricant Eye Ointment; Refresh) were applied. Next a vasculature image was obtained under green light, then the hemodynamic response to a visual stimulus was imaged under red light. The stimulus consisted of an alternating checkerboard pattern (20° wide bar, 25° square size) moving across a mean luminance gray background. On each trial, the stimulus bar was swept across the four cardinal axes 10 times in each direction at a rate of 0.1 Hz. Up to 10 trials were performed on each mouse. A minimum of three trials were averaged to produce altitude and azimuth phase maps, calculated from the discrete Fourier transform of each pixel. A “sign map” was produced from the phase maps by taking the sine of the angle between the altitude and azimuth map gradients. In the sign maps, each cortical visual area appears as a contiguous red or blue region (Figure 3c).

### Magnetic resonance imaging

P64-P151 mice underwent either the SHIELD surgical protocol as described here or our existing 5mm window protocol, with the following modifications: a headframe made of yttria partially stabilized zirconia was used instead of a titanium one, and the well was replaced with a custom 3D printed cap.

100 µm isometric T1-weighted anatomical scans were taken of each mouse 8-26 days after the initial surgery. 24 hours before the MRI scan, mice received an IP injection of 100 mM MnCl2 and 100 mM bicine scaled to achieve a dose of 66 mg of manganese per kg (Massad and Paulter, 2011). Mice were anesthetized using isoflurane and T1-weighted anatomical scans were taken using a RARE pulse sequence (Henning et al 1986).

Because the headframes are cemented to the skull after being stereotaxically aligned to bregma, we used the position of the headframe in our anatomical scans to infer the location of bregma. As the zirconia headframes are not visible in T1-weighted MRI volumes, we coated the headframes with white-petroleum-based eye lubricant (Systane nighttime eye ointment) to visualize the outer surface of the headframe. SHIELD-zirconia headframes had additional fiducial markers filled with white-petroleum to facilitate this alignment.

To analyze the deformation caused by the surgical implant, we identified voxels belonging to the brain with a UNET (Hsu L-M. et al. 2021) fine-tuned using data collected under the same imaging protocol. These segmentations were manually corrected using 3D Slicer (v4.11) (Fedorov A. et al 2012). Surgically naïve brains from p65 mice (Qiu et al. 2018) were manually segmented with 3D Slicer. We then used 3D Slicer to generate a mesh of each brain surface that was aligned to bregma in the anterior-posterior axis, rotated so that bregma and lambda were level using the skull if possible or the headframe if the skull was not visible, and aligned to the midline of the brain in the medial-lateral axis. We found where the mesh crossed each of ten coronal planes along the anterior-posterior axis. The right surgically naïve hemisphere was mirrored and compared to the left hemisphere, and the area between these two surfaces was computed using triangulation.

### Skull contour mapping

Mice were prepared for surgery and anesthetized as described above. A headplate was installed, and the scalp was removed to reveal the skull. The mouse was then euthanized and scans were acquired ex vivo with a Micro Epsilon Laser Line Scanner. The point clouds from each scan were converted to a surface using Geomagic Design X. All skull surfaces were then imported into Solidworks.

### Implant preparation prior to Neuropixels recordings

#### SORTA-Clear plug removal and agarose application

To prepare the brain for recording, the SORTA-Clear coating over the implant is removed and replaced with a temporary layer of Kwik-Cast (World Precision Instruments). The mouse is anesthetized with isoflurane (5% induction, 1-2% maintenance, 100% O2) and eyes protected with ocular lubricant (IDrop, VetPLUS). Body temperature is maintained at 37.5°C (TC-1000 temperature controller, CWE, Incorporated). The well is cleaned of any debris using ethanol swabs. Then, the inside of the well surrounding the SORTA-Clear plug is painted with white Metabond to improve visibility during probe insertion. Once the Metabond is dry, the well is flooded with enough ACSF to completely submerge the SORTA-Clear sheet, which is then removed with small forceps, starting at the anterior or posterior end of the sheet and peeling gently to remove it in one piece. Once the SORTA-Clear sheet is completely detached from the implant, the edges of implant holes are tested with small forceps to ensure all holes are free of SORTA-clear or debris. ACSF is removed from the well using Sugi spears (Kettenbach) and the well is filled with Kwik-Cast. Once the Kwik-Cast is fully dry, a plastic protective cap is secured on the well to protect against debris and the mice are returned to their home cage. This preparation is stable for at least three days before the recording.

### Neuropixels recordings

#### Preparation of mouse for recording

The mouse is removed from its home cage and clamped to the running wheel on the experimental rig. Wheel height is adjusted as needed for each mouse. Once head-fixed, the protective well cap and Kwik-Cast layer are removed. The ground wire is tucked into the side of the well and any excess debris cleaned using a Sugi spear or cotton tipped applicator. Approximately 0.4 ml of agarose is applied in a smooth layer over the entire implant surface and ground wire. After popping any large bubbles, the agarose is allowed to set for 10 seconds. To prevent the agarose from drying out during the experiment, a layer of silicon oil is applied over exposed agarose with a toothpick.

For the optotagging experiment, Dura-Gel (DOWSIL™ 3-4680) was used in the place of agar. Dura-gel was applied directly after removing the SORTA-Clear plug. We found that mixing the white and blue components at a 35:50 ratio produced a tougher consistency that was less likely to coat the probes in residue. After stirring for several minutes to ensure a homogenous mixture, the head of the mouse was tilted to level the well, the ground wire was placed so that it rested on the brain surface, and Dura-Gel was applied a few drops at a time with the broken end of a cotton swab. Any bubbles that formed in the holes of the implant were pushed out with the applicator. Dura-gel was applied until the layer was at least 1 mm thick and then left to cure for at least 10 minutes, after which the mouse was returned to its home cage. The Dura-Gel was given at least 24 hours to fully cure before recording.

#### Application of CM-DiI and DiO to probes

In our experiments, probes were inserted into the same mouse on sequential days. To distinguish the paths of these different penetrations, we used two dyes, CM-DiI (1 mM in ethanol; ThermoFisher Product #V22888) and DiO (1mM in DMF; ThermoFisher Product #V22886), on separate recording days. The probes were coated with dye before recordings by immersing them at least 3mm into a well filled with dye. Each probe was dipped five times to ensure adequate coating.

#### Probe insertion

Our custom experimental rig can insert up to six Neuropixels probes simultaneously (Durand et al. 2023). Each probe is mounted on a separate 3-axis micromanipulator with a 15 mm travel range (New Scale Technologies, Victor, NY). Probe and implant hole combinations were selected prior to each experiment. Probes were driven to their target holes and lowered to the surface of the brain while the operator monitored a camera feed to avoid vasculature and watched real-time signals on the OpenEphys GUI to identify activity indicative of the brain surface. If the probe needed adjustment when attempting to insert (e.g. to avoid vessels), the probe was completely retracted out of the silicon oil to prevent probe bending. If a probe could not be inserted into its assigned hole, another target hole was selected. Once all probes reached the brain surface, each probe was zeroed and set to insert 3000 µm at 200 µm/min. Insertion of a probe was stopped if its base bumped that of other probes (when probes target nearby locations, care must be taken to avoid collisions between the probe bases). Once all probes reached their final depth, the probes were allowed to settle for 10 minutes, and photo documentation of the inserted probes was captured. In a subset (23) of recordings, probes were inserted to 3100 µm then retracted 100 µmm to their final depths to reduce tissue compression and subsequent electrode drift relative to the brain (Figure S4).

Our goal was to insert 6 probes per session. Overall, we achieved a penetration success of 5.46 probes per session, with failures due to dura regrowth, collisions with other probes, or probe breakage during manipulation. To facilitate insertion through dura regrowth, the tips of two of six Neuropixels probes were sharpened as previously described (Durand et al. 2023), which we found to effectively reduce insertion failures (Figure S4).

#### Probes and grounding

All neural recordings were carried out with Neuropixels 1.0 probes, as previously described (Durand et al. 2023; Siegle et al. 2021). The 383 electrodes closest to the tip were used, providing a maximum of 3.84 mm of tissue coverage. The signals from each recording site are split in hardware into a spike band (30 kHz sampling rate, 500 Hz highpass filter) and an LFP band (2.5 kHz sampling rate, 1000 Hz lowpass filter).

A 32 AWG silver wire (A-M Systems) is epoxied to the headframe during the implant surgery and serves as the ground connection. The wire is pre-soldered to a gold pin embedded in the headframe well, which mates with a second gold pin on the protective cone. This second gold pin is connected to both the behavior stage and the probe ground. Prior to the experiment, the brain-to-probe ground path was checked using a multimeter. The reference connection on the Neuropixels probes was permanently soldered to ground using a silver wire, and all recordings were made using the tip reference configuration. The headstage grounds (which are contiguous with the Neuropixels probe grounds) were connected with 36 AWG copper wire (Phoenix Wire). All probes were connected in parallel to animal ground.

#### Data acquisition

Neuropixels data was acquired at 30 kHz (spike band) and 2.5 kHz (LFP band) using the Open Ephys GUI (Siegle et al. 2017). Gain settings of 500x and 250x were used for the spike band and LFP band, respectively.

Videos of the eye, body, and face were acquired at 60 Hz. The angular velocity of the running wheel was recorded at the time of each stimulus frame, at approximately 60 Hz. Synchronization signals for each frame were acquired by a dedicated computer with a National Instruments card acquiring digital inputs at 100 kHz, which was considered the master clock. A 32-bit digital “barcode” was sent with an Arduino Uno (SparkFun DEV-11021) every 30 s to synchronize all devices with the neural data.

To synchronize the visual stimulus to the master clock, a silicon photodiode (PDA36A, Thorlabs) was placed on the stimulus monitor above a “sync square” that flipped from black to white every 60 frames.

#### Optogenetic silencing of cortex

To silence cortex, a 488 nm blue laser spot ( 100 µm diameter) was steered over dorsal cortex with a galvo system in a VGAT-ChR2-EYFP mouse expressing channelrhodopsin-2 (ChR2) in GABAergic neurons (JAX strain #014548). A Neuropixels probe was inserted into VISp to monitor cortical activity while the laser power and position was varied (three powers: 1, 2 and 12 mW; four positions relative to recording location: 0, 0.5, 1 and 2 mm anterior). The laser stimulus was timed to coincide with a visual stimulus to drive visual cortical activity (windowed vertical grating: 500 ms duration, 50 degree diameter, 0.04 cycles per degree). Only regular spiking units were included in analysis, defined by a peak-to-trough duration of greater than 0.4 ms measured on the peak channel waveform.

#### Optotagging in the midbrain

To perform optotagging in the midbrain, AAV2-hSyn-FLEX-ChrimsonR-tdTomato (UNC Vector Core) was injected into the left midbrain reticular nucleus (MRN) of Vglut2-Cre mice (JAX strain #028863). One 200 nL injection was done at 3.8 mm posterior to bregma, 0.87 mm from midline, and at a depth of 3.3 mm from pia. The virus was given 4 weeks to express before electrophysiological recordings. At that time, the mouse was head-fixed on a running wheel and a recording was performed with a Neuropixels probe inserted 4500 um deep.

Vglut2+ cells were identified by evaluating neural responses to five pulses of red light (638 nm), presented at a rate of 20 Hz and with each pulse lasting 10 ms. Light was delivered via a collimator (Thorlabs CFC2-A) sending a beam of laser light about 400 um wide to the surface of the SHIELD implant, near the insertion location of the Neuropixels probe. Trials with different laser powers were randomly interleaved, with a randomized intertrial interval in the range from 0.8 to 1.2 seconds. 50 trials were performed at each power. Vglut2+ cells were identified by their significant (p < 0.05) increase in firing rate during all five laser pulses.

#### Probe removal and cleaning

At the end of the recording sessions probes were removed slowly from the brain (approximately 1 mm/minute). A Sugi spear was used to clear off excess silicone oil from the implant and forceps were used to remove the agarose. The well was filled with Kwik-Cast, allowed to dry for 30 seconds, and then the well cap was replaced.

To remove debris, probes were submerged in a 1% Tergazyme solution overnight then rinsed in deionized water the next day.

### Visual stimulation and behavior

Mice were trained to perform a visual change detection task in which one of 8 natural images was continuously flashed (250 ms image presentation followed by 500 ms gray screen) and mice were rewarded for licking when the image identity changed. The change detection task is described in previous publications (Garrett et al. 2020; Groblewski, Ollerenshaw, et al. 2020).

Receptive fields were mapped during each experiment using Gabor stimuli windowed to have a 20 degree diameter and presented in a 9x9 grid. Grid positions were spaced by 10 degrees. Gabors had a temporal frequency of 4 Hz, a spatial frequency of 0.08 cycles/degree and were shown at 3 orientations (0, 45, 90 degrees). They were 250 ms in duration without gaps between stimuli. There were 15 trials for each condition (81 positions, 3 orientations).

### Ex vivo imaging and analysis to identify probe tracts

#### TissueCyte imaging

Serial two-photon tomography was used to obtain a three-dimensional (3D) image volume of coronal brain images for each specimen. This 3D volume enables spatial registration of each mouse’s brain to the Allen Mouse Common Coordinate Framework (CCF). Methods for this procedure have been described in detail in whitepapers associated with the Allen Mouse Brain Connectivity Atlas and in Oh et al (Oh et al. 2014).

#### Probe track annotation and CCF Registration

Probe tracks were annotated manually in the TissueCyte image volume after resampling to 10x10x10 µm voxels. Because linear probe trajectories become curved after warping to the CCF, annotations were performed before applying local deformation to the volume, enabling us to fit a line to the annotated track points for each probe. Electrode positions were adjusted along this line based on electrophysiological features and unit density. After this refinement process, electrode coordinates were passed through the final deformation field and thus warped into the CCF.

For probe insertions into visual cortical areas, vasculature landmarks were used to warp probe insertion images to corresponding ISI images which delineated the visual area boundaries for each animal. Probe insertions were then annotated on these warped images and manually assigned to visual cortical areas based on ISI-derived visual sign maps. These manual area assignments superseded CCF area labels for cortical units.

### Histology

Brains were perfused with 4% paraformaldehyde (PFA), post fixed in 4% PFA overnight, then stored in 30% sucrose solution in preparation for sectioning. Brains were sectioned at 50 µm on a freezing microtome. Rinsed free floating sections were blocked in a solution containing 5% donkey serum for one hour. They were then incubated for 24 hours at room temperature with 1:1000 Mouse anti-GFAP and 1:1000 Rabbit anti-NeuN in blocking solution. Following a sequence of rinses, sections were incubated in 1:500 donkey anti-mouse 568, 1:500 donkey anti-rabbit 488, and 1:1000 DAPI in blocking solution. After final rinses, sections were mounted on slides and imaged with an Olympus VS120 fluorescent slide scanning system.

### Data pre-processing and spike sorting

Prior to spike sorting, the spike-band data passed through 4 steps: offset removal, median subtraction, filtering, and whitening. First, the median value of each channel was subtracted to center the signals around zero. Next, the median across channels was subtracted to remove common-mode noise. To remove noise sources that are shared across channels, the median was calculated across channels that are sampled simultaneously as previously described (Siegle et al. 2021). The median-subtracted data was sent to Kilosort2, which applies a 150 Hz high-pass filter, followed by whitening in blocks of 32 channels. The filtered, whitened data was saved to a separate file for the spike sorting step.

Kilosort2 was used to identify spike times and assign spikes to individual units (Pachitariu et al. 2023). The Kilosort2 algorithm will occasionally fit a template to the residual left behind after another template has been subtracted from the original data, resulting in double-counted spikes. This can create the appearance of an artificially high number of inter-spike interval violations for one unit or artificially high zero-time-lag synchrony between nearby units. To eliminate the possibility that this artificial synchrony would contaminate data analysis, the outputs of Kilosort2 was post-processed to remove spikes with peak times within 5 samples (0.16 ms) and peak waveforms within 5 channels ( 50 µm).

### Data analysis

#### Quality metric criteria for unit inclusion

For computing quality control metrics and reporting the number of units recorded in Figure 4, all units were included except those with non-physiological waveforms. Briefly, units with abnormal spatial spread (>25 channels), abnormal temporal waveform (absence of peak and trough) or multiple spatial peaks were deemed to be noise as described previously (Siegle et al. 2021). For all other analysis, units were included if they met our standard quality control criteria: inter-spike violation rate less than 0.5, amplitude cutoff less than 0.1, and presence ratio greater than 0.95. Each of these metrics was calculated as described previously (Siegle et al. 2021).

#### Identification of alpha events

Alpha events in VISp and MOp were identified with the following algorithm, which was empirically found to agree with manual annotation. First, unit activity within each area was binned into 10 ms bins and averaged together to create a population spike rate, which was then smoothed with a symmetrical exponential filter (10 ms tau). Sessions with less than 20 neurons in the relevant area (VISp or MOp) were excluded in this analysis. Rhythmic pauses in activity in the resulting population trace were then identified by finding local maxima in the inverted trace (scipy.signal.find_peaks; prominence=5, height=-1, width=[100 ms, 300 ms]). An alpha event was registered if there were at least four maxima at intervals between 125 and 350 ms in a 1.25 second window. After an alpha event, we imposed a refractory period of two seconds before another event could be registered. Because this algorithm occasionally identified non-rhythmic activity, we further filtered the resulting events by requiring that the power of the autocorrelogram in the 3-7 Hz band exceed 0.1 (spikes/sec).

#### Cross-correlograms and coherence

For each session, cross-correlograms (CCGs) were computed between the population activity of VISp or MOp and all other regions for which at least 20 neurons were recorded. CCGs were computed for 2 second windows centered on the alpha event and were normalized so that the auto-correlogram had a peak of 1. CCGs for each area pair were then pooled across sessions for subsequent analysis.

Multi-taper coherence was computed using the nipy software package (https://github.com/nipy/nitime) (Millman & Brett 2007) for four second windows centered on each alpha event. To isolate coherence due to alpha rhythms, coherence was also calculated for random times during each session and subtracted from coherence calculated during alpha events.

#### Receptive Field Mapping

The receptive field for one unit was defined as the 2D histogram of spike counts at each of 81 locations of the Gabor stimulus (9 x 9 matrix). The significance of a given unit’s receptive field was measured with a chi-square test, as described previously (Siegle et al. 2021). Any unit with a p-value less than 0.01 was considered to have a significant receptive field.

#### Statistics

Unless otherwise specified, paired comparisons were made with the Wilcoxon signed-rank test and unpaired comparisons with the Wilcoxon rank-sums test. For panels with multiple comparisons, p-values were corrected with the Benjamini-Hochberg method.

## Data and code availability

Data and code used in this publication will be available via online repositories at the time of publication. CAD models for all parts are available for download here: https://portal.brain-map.org/explore/toolkit/hardware.

## Acknowledgments

We thank the Allen Institute founder, Paul G. Allen, for his vision, encouragement and support. Funding for this project was provided by the Allen Institute. This project was initiated as part of the MindScope Program at the Allen Institute; we thank the leadership of MindScope, Christof Koch and John Phillips, for their support and advice. We thank Karel Svoboda for advice on the manuscript. We thank Kat North and Jerome Lecoq for advice on surgical techniques. We thank the Transgenic Colony Management, Animal Care, and Laboratory Animal Services teams for caring for the mice in this study. We thank the Imaging team for performing TissueCyte imaging. We thank Suhasa Kodandaramaiah and members of his laboratory for discussion and advice regarding methods associated with their “See-Shells” method.

## Author contribution matrix

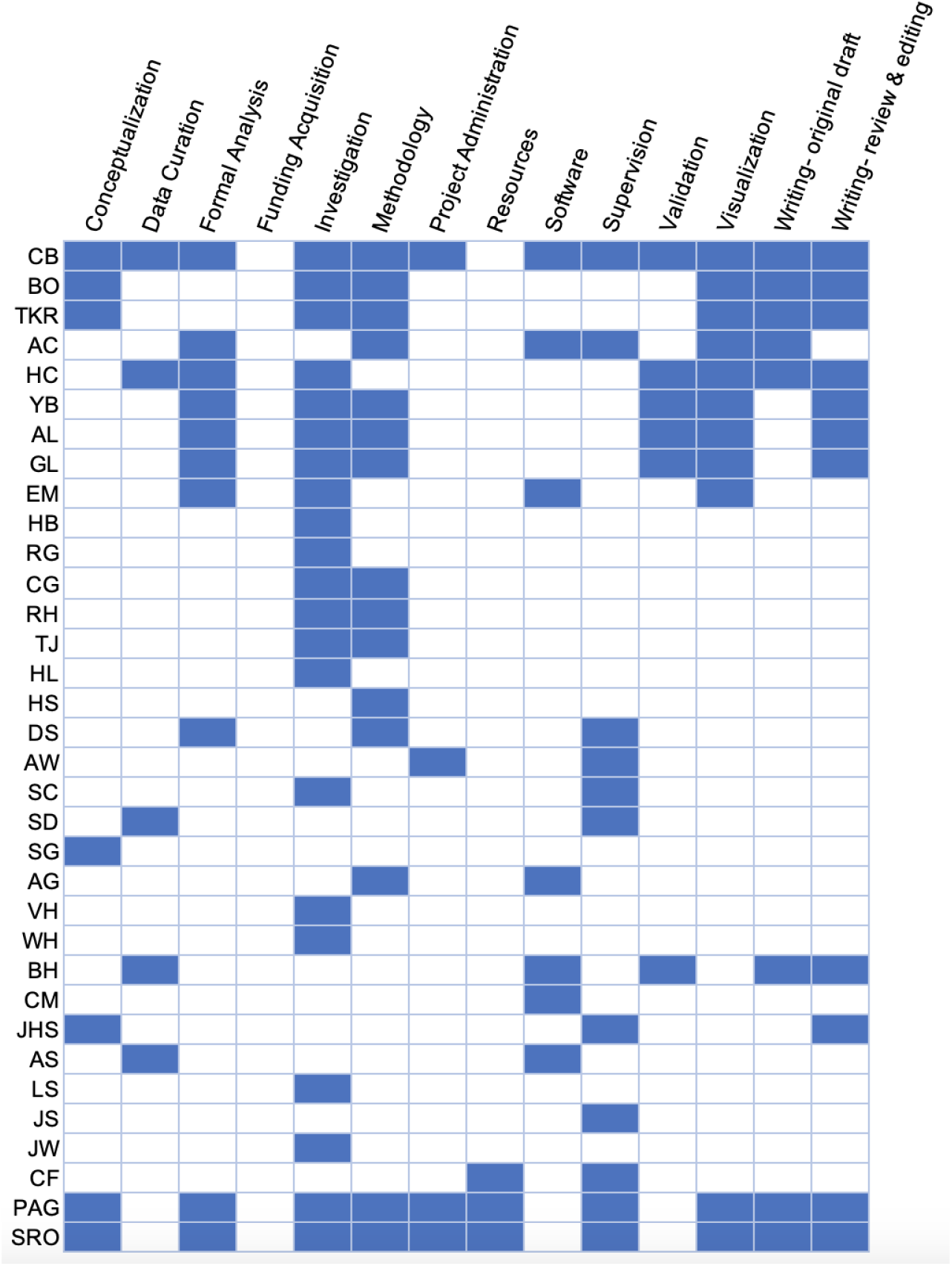

---

**Figure S1:**
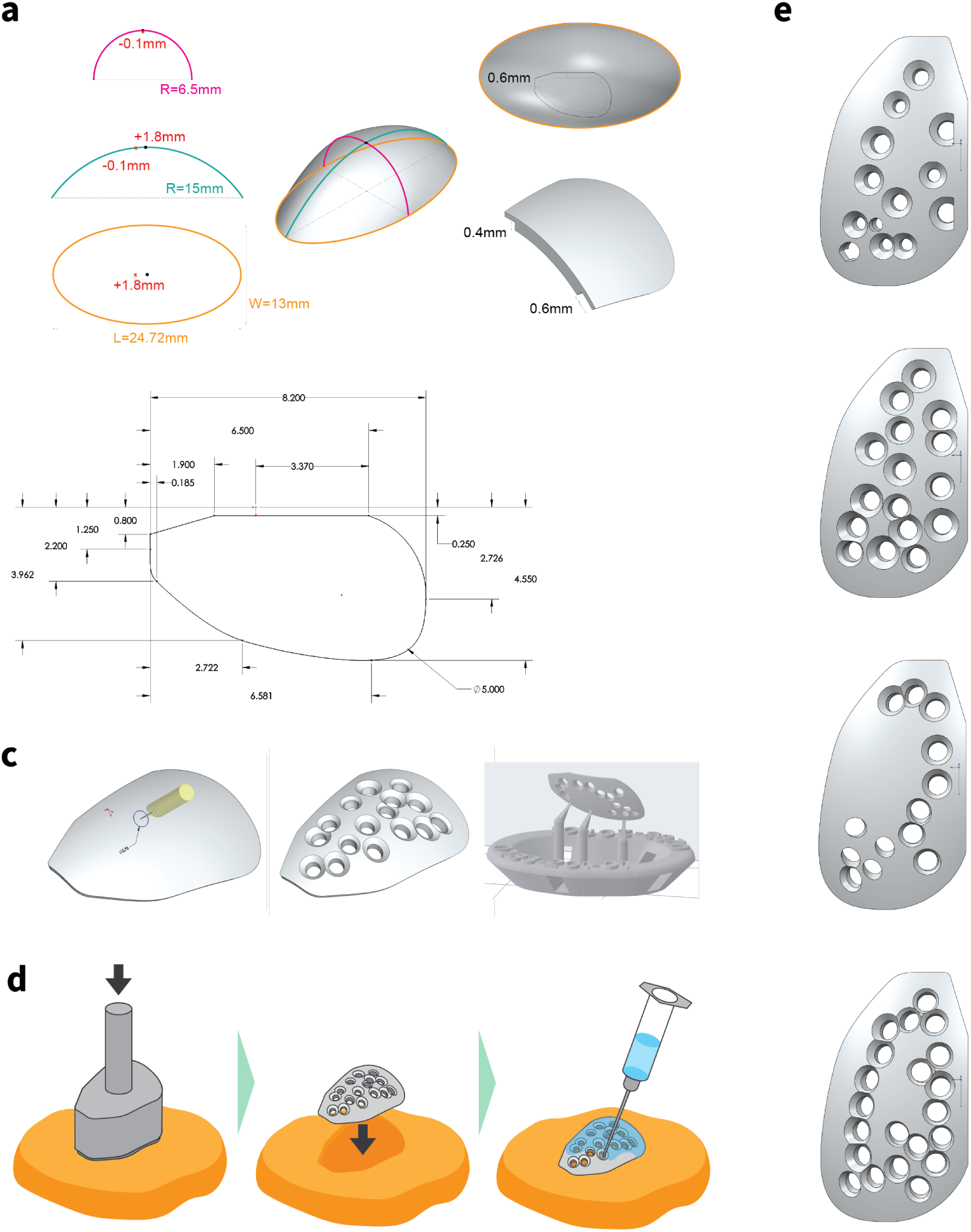
Implant creation and coating. **a,** Using computer-aided design (CAD) software, the implant design was initially created as a triaxial ellipsoid-like surface intended to match the curvature of an average mouse skull in the area where the craniotomy is located (see Figure 2a). The surface is defined by three curves with respect to Bregma (red x). The shape of the implant is cut out of the ellipsoid surface. The flange extends 0.6 mm beyond the craniotomy, providing a surface to glue the implant to the skull. The inner portion circumscribed by this flange extends down into the craniotomy 0.4 mm and sits directly on the brain. **b,** Top-down (X and Y) dimensions (in mm) of the implant. **c,** Implant holes are created over target coordinates at an angle that matches Neuropixels probe insertion angles. Each hole receives 45-degree chamfers, allowing for some flexibility in probe insertion angles. Implants are then 3D-printed in clear resin at the orientation shown, UV cured at 60C for 15min, and rinsed with isopropyl alcohol prior to storage. **d,** Coating implants with SORTA-Clear requires making an impression of the implant in putty using a custom stamp tool. The implant is then gently placed into the impression. SORTA-Clear is loaded into a 1mL syringe with 25g needle and each hole is slowly filled to prevent air from getting trapped. After filling each hole, the entire surface of the implant is covered with a thin layer leaving the lip portion at the edges uncovered. Coated implants are allowed to cure for 24h before being removed from putty. They are then rinsed with 70% EtOH, dried, and individually packaged for the autoclave. **e,** The four SHIELD implants, each with unique probe hole configurations, used in this study.

**Figure S2:**
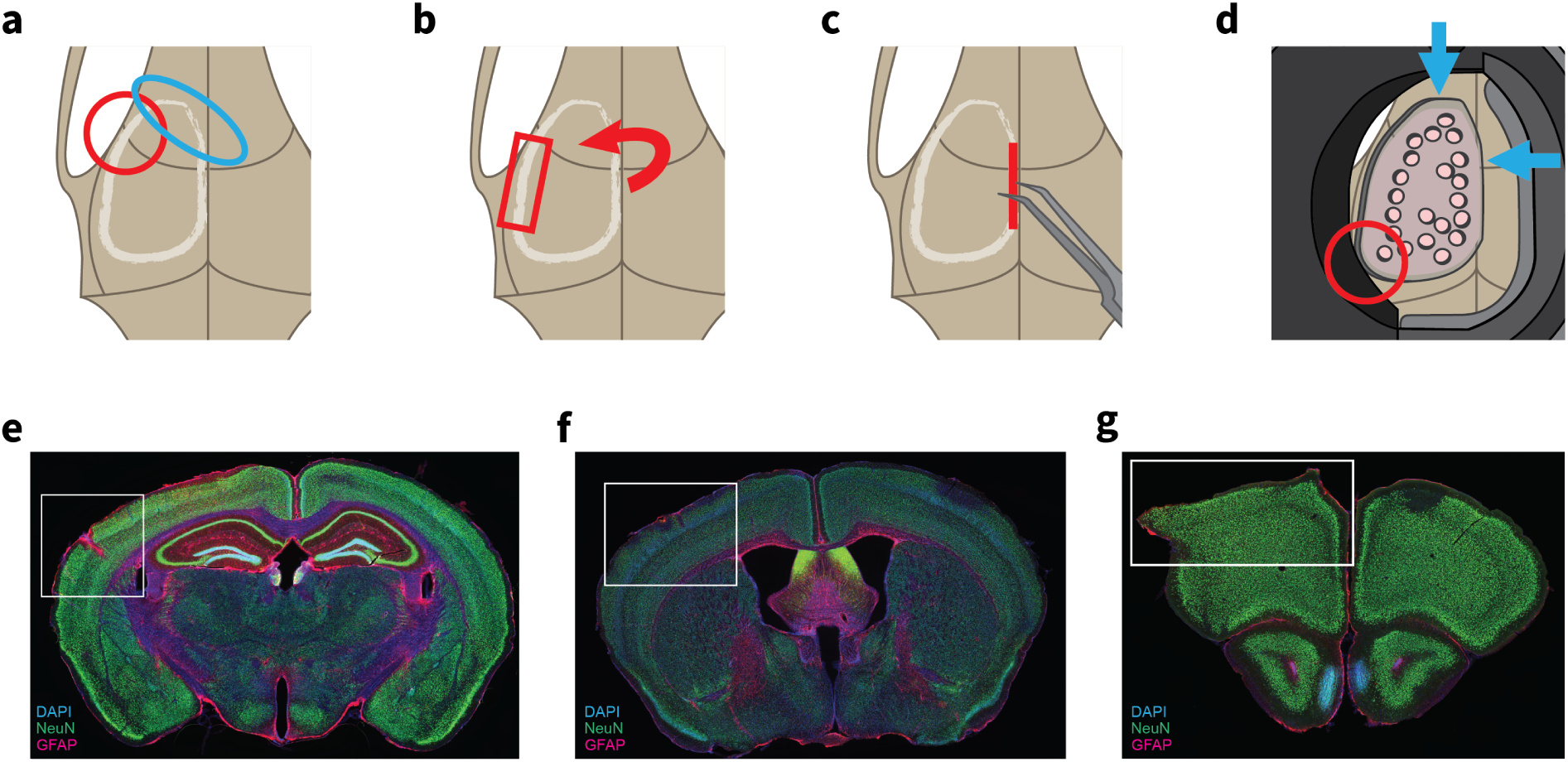
Surgical tips and troubleshooting. **a,** The drilling depth in the upper left corner of the craniotomy can vary substantially due to small variations in proximity to the eye socket (red). The drill path comes very close to the rostral rhinal vein and heavy vasculature near its intersection with the sagittal sinus (blue). **b,** While it is advised to drill until a crack or separation occurs over most of the drill path, it is often helpful to leave a very thin layer of bone on the lateral edge (red rectangle) of the craniotomy to act as a hinge when removing the skull. **c,** Removal of the drilled skull should begin from the medial side. **d,** When placing the implant, align it to the anterior and medial edges (blue arrows) to ensure accurate placement, as the lateral edge is more difficult to visualize. Clearance between the well, metabond, and implant can be very tight in the lower lefthand corner, so care should be taken to keep the metabond low enough to account for the implant curvature of the implant lip (red circle). **e-g,**. Histological examples of tissue damage caused excessive drilling (e), dura tearing during durotomy (f), and improper seating of the implant in the craniotomy (g).

**Figure S3:**
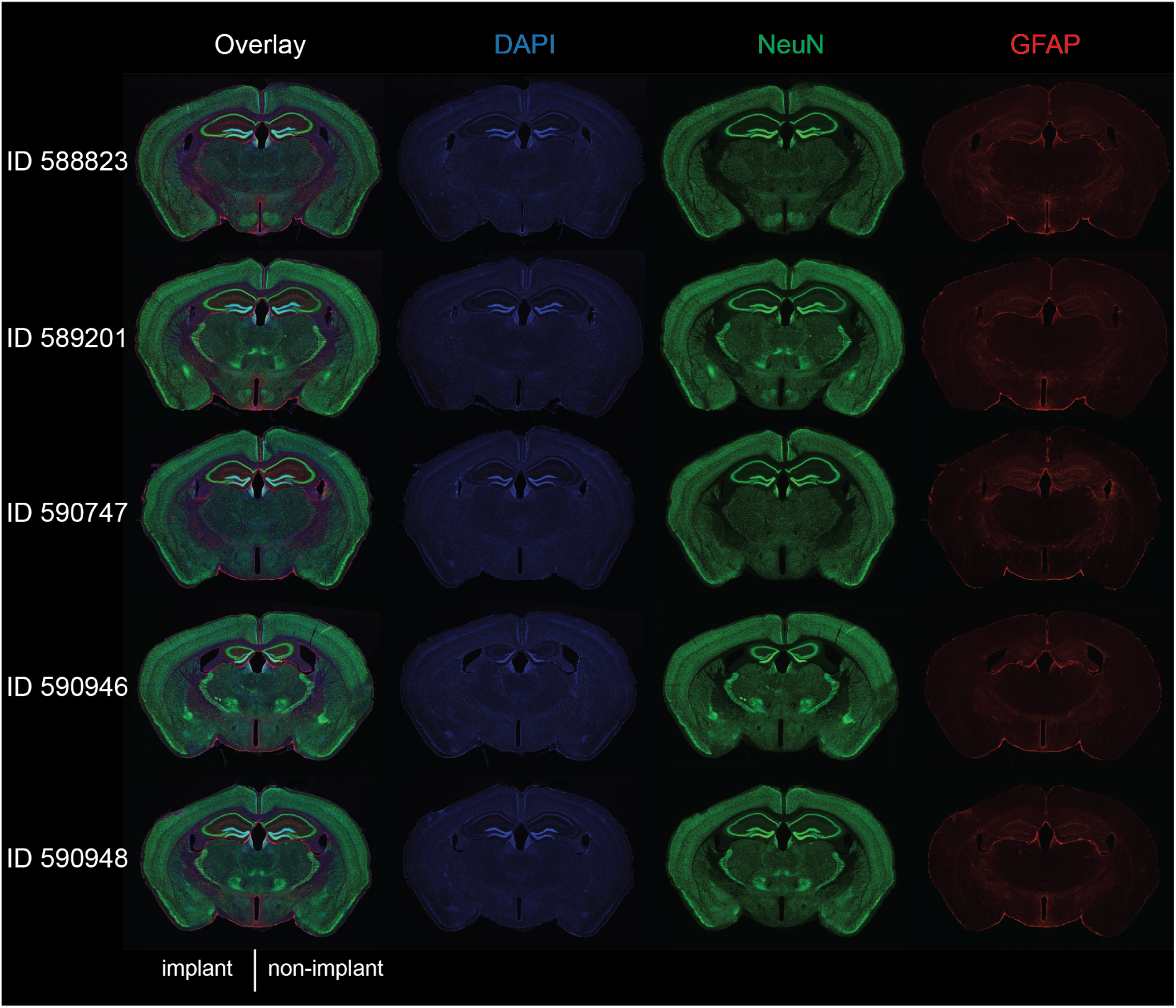
Additional histological examples from five mice. SHIELD implant was installed over the left hemisphere.

**Figure S4:**
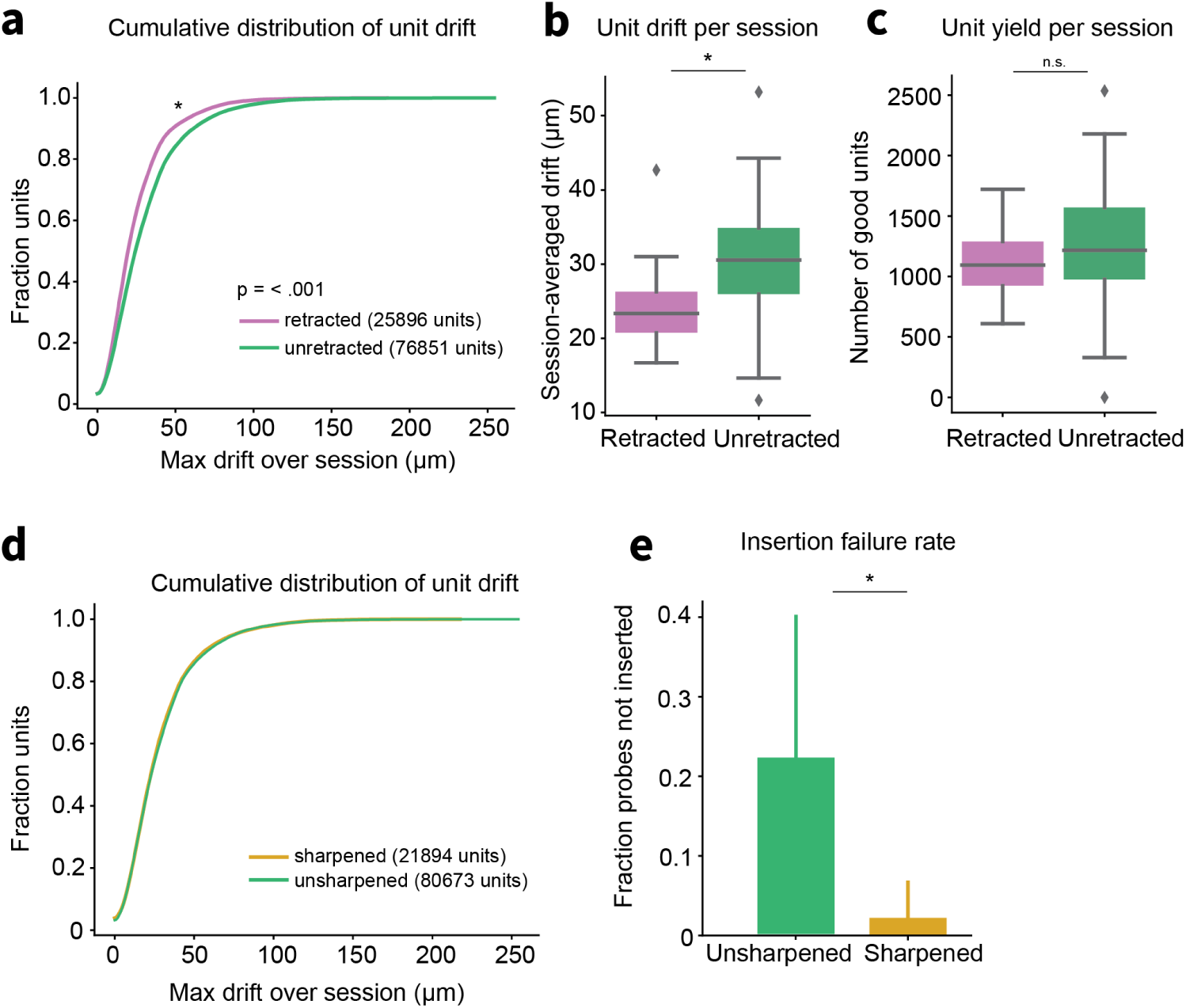
Neuropixels probe insertion optimization. **a,** Retracting Neuropixels probes 100 µm after inserting reduces unit drift. Cumulative distribution of point-to-point drift per unit over session with (pink, n = 25,896 units) and without (green, n = 76,851 units) probe retraction. Probes were retracted on a session-by-session basis. Units from sessions where probes were retracted had lower drift (mean = 24.48 µm) than those from sessions where probes were not retracted (mean = 30.25 µm; Kolmogrov-Smirnov test, p < .001). **b,** Sessions in which Neuropixels probes were retracted show less drift. Average drift across all units recorded in retracted sessions (n = 23, mean = 24.23 µm) was less than non-retracted sessions (n = 60, mean = 30.56 µm). **c,** Number of good units recorded per session is similar across retracted and non-retracted sessions. Histogram showing distribution of unit yield for retracted (mean = 1125 units) and non-retracted (mean = 1280 units) sessions. Distributions did not differ significantly (Kolmogrov-Smirnov test, p = 0.318). **d,** Sharpened Neuropixels probes show comparable drift to unsharpened probes. Cumulative distribution of point-to-point drift over session per unit with (yellow, n = 21,894 units) and without (green, n = 80,673) sharpened probe tips. **e,** We sharpened 2 of 6 Neuropixels probe tips during data collection. Insertion failure rate for those probes before sharpening (green, n = 32 insertions) and after sharpening (gold, n = 112) is shown with 95% confidence intervals (binomial test). Sharpened Neuropixels probes have a lower insertion fail rate than unsharpened probes (p = 0.0004, Fisher’s exact test).

**Figure S5:**
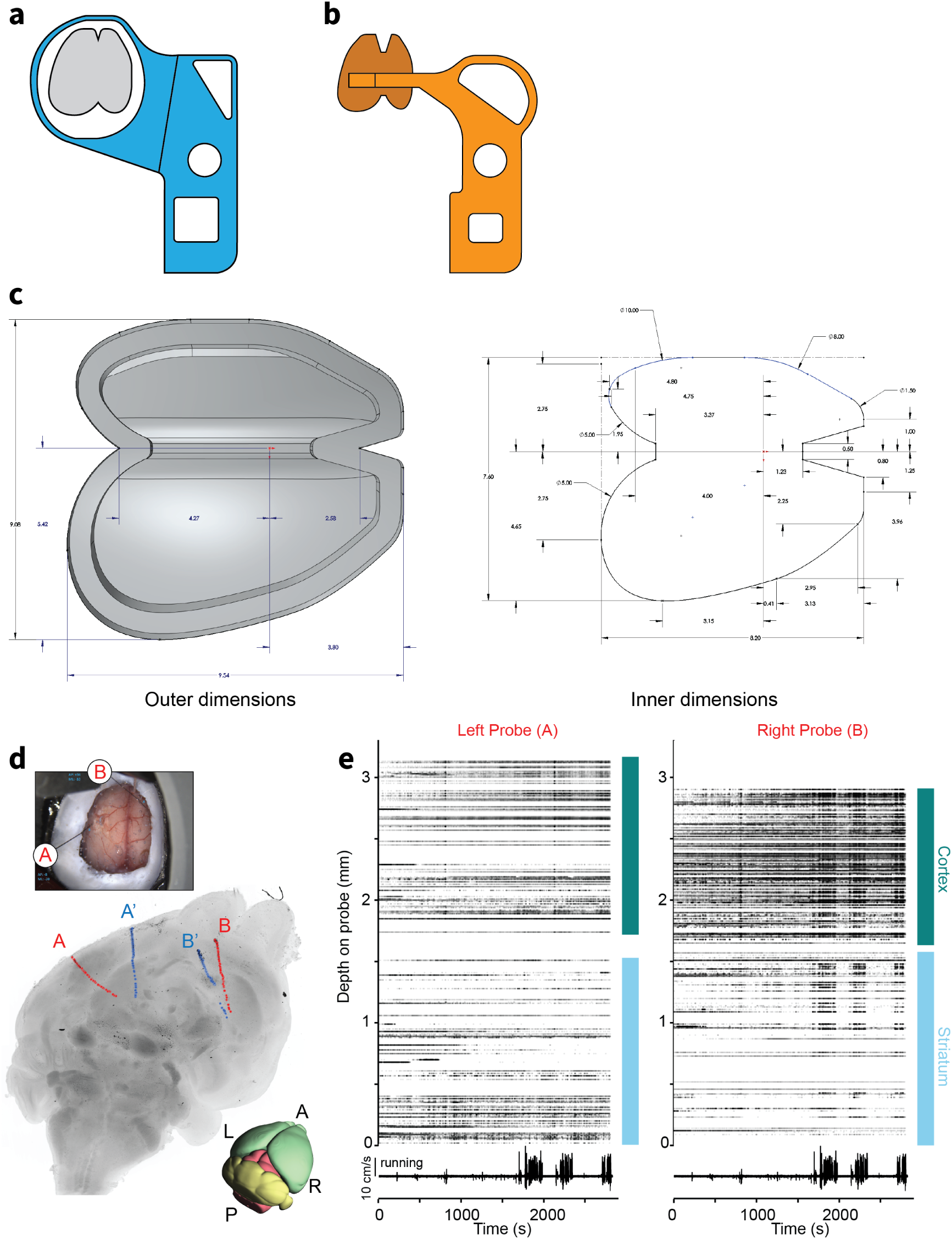
Bilateral SHIELD implant and example recording. **a,** Headframe used with bilateral implant. **b,** Tracer for bilateral craniotomy. **c,** Implant dimensions. **d,** Top, Neuropixels insertions into right and left brain hemispheres. Bottom, reconstruction of probe tracts from two experimental sessions with bilateral insertions. Blue and red indicates recordings performed on separate days. **e,** Raster plot of spiking activity recorded from units across each probe. Insertions spanned cortex and striatum. Marker intensity indicates spike amplitude.

**Figure S6:**
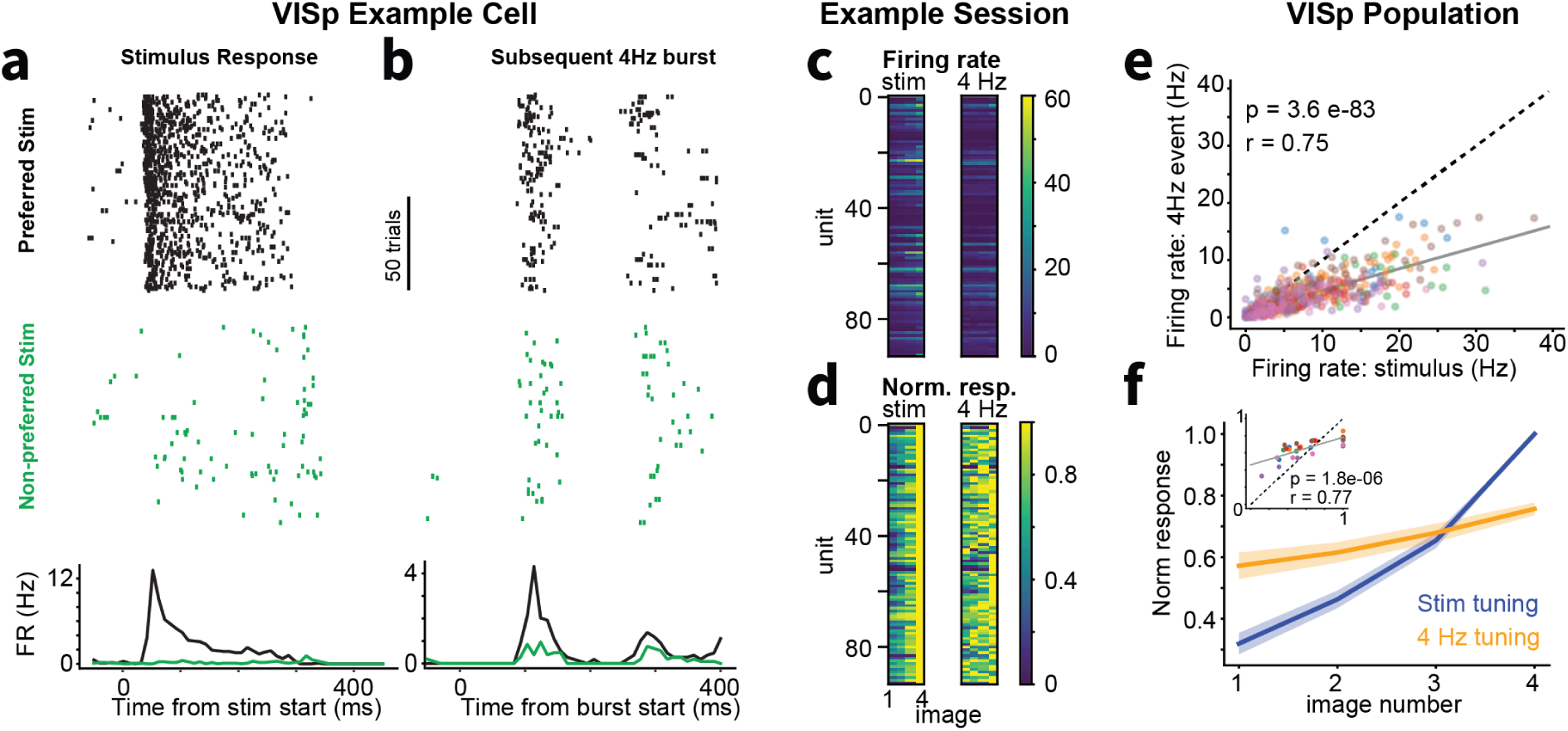
Replay of recent sensory activation during alpha events in VISp. **a,** Spike rasters for one example VISp unit during presentation of a preferred natural image (black) and a non-preferred image (green). (Bottom) PSTH of same unit for preferred and non-preferred image presentations. The image was presented for 250 ms starting at time zero. **c,** Rasters for same unit during alpha burst events following stimulus presentation trials in (a). Note that this unit responds more vigorously following presentation of its preferred stimulus. **c,** (Left) Heatmap showing tuning of all VISp units recorded during an example experiment for the four stimuli presented during the session. Rows depict the mean response for individual units to each of the four stimuli shown during this session sorted by magnitude. (Right) Heatmap showing response of same units to the alpha burst event following visual stimulation. **d,** Same as (c) but row normalized. **e,** Correlation of firing rate during visual stimulation and subsequent alpha burst. Each dot is a VISp unit, color-coded by recording session. **f,** Mean stimulus tuning across all VISp units for the sensory-evoked response (blue) and the subsequent alpha burst (orange). This plot is the column average of (d) but across all recording sessions. Inset: Correlation of 4Hz tuning curve and stimulus tuning curve across sessions. Points are colored by session. Each session contributes four points to the plot.

